# PAX4 loss of function alters human endocrine cell development and influences diabetes risk

**DOI:** 10.1101/2022.05.15.491987

**Authors:** Hwee Hui Lau, Nicole A. J. Krentz, Fernando Abaitua, Marta Perez-Alcantara, Jun-Wei Chan, Jila Ajeian, Soumita Ghosh, Benoite Champon, Han Sun, Alokkumar Jha, Shawn Hoon, Nguan Soon Tan, Daphne Gardner, Shih Ling Kao, E Shyong Tai, Anna L Gloyn, Adrian Kee Keong Teo

**Author notes:** **Correspondence:** Adrian Teo, Ph.D., Institute of Molecular and Cell Biology (IMCB), A*STAR, Department of Biochemistry and Department of Medicine, Yong Loo Lin School of Medicine, NUS, Anna L. Gloyn, DPhil, Centre for Academic Medicine, Division of Endocrinology & Diabetes MC 5660, Department of Pediatrics, Stanford School of Medicine, Stanford University, 453 Quarry Road, Palo Alto, CA 94304-5660. These authors jointly contributed to this work (Hwee Hui Lau, Nicole A. J. Krentz). These authors jointly supervised this work (Anna L Gloyn, Adrian Kee Keong Teo).

## Abstract

Diabetes is a major chronic disease with an excessive healthcare burden on society^1^. A coding variant (p.Arg192His) in the transcription factor *PAX4* is uniquely and reproducibly associated with an altered risk for type 2 diabetes (T2D) in East Asian populations^2–7^, whilst rare *PAX4* alleles have been proposed to cause monogenic diabetes^8^. In mice, *Pax4* is essential for beta cell formation but neither the role of diabetes-associated variants in *PAX4* nor PAX4 itself on human beta cell development and/or function are known. Here, we demonstrate that non-diabetic carriers of either the *PAX4* p.Arg192His or a newly identified p.Tyr186X allele exhibit decreased pancreatic beta cell function. In the human beta cell model, EndoC-βH1, *PAX4* knockdown led to impaired insulin secretion, reduced total insulin content, and altered hormone gene expression. Deletion of *PAX4* in isogenic human induced pluripotent stem cell (hiPSC)-derived beta-like cells resulted in derepression of alpha cell gene expression whilst *in vitro* differentiation of hiPSCs from carriers of *PAX4* p.His192 and p.X186 alleles exhibited increased polyhormonal endocrine cell formation and reduced insulin content. *In silico* and *in vitro* studies showed that these *PAX4* alleles cause either reduced PAX4 expression or function. Correction of the diabetes-associated *PAX4* alleles reversed these phenotypic changes. Together, we demonstrate the role of PAX4 in human endocrine cell development, beta cell function, and its contribution to T2D-risk.

## Introduction

Diabetes is a chronic condition affecting more than 537 million people worldwide, giving rise to devastating complications and healthcare burdens on society^1^. In an effort to identify novel disease-causing mechanisms and tractable targets for therapeutic development, numerous genome-wide studies have been performed across different ancestries to identify genetic variants that influence diabetes risk^2–4^. An Asian-enriched *PAX4* p.Arg192His (rs2233580) coding variant has been reproducibly associated with T2D risk [odds ratio of ∼1.75]^2,5^. Additional studies revealed that 21.4% of 2,886 people with early-onset T2D carried at least one *PAX4* p.Arg192His allele^6^. Carriers of the T allele (p.His192) have a dose-dependent earlier age of T2D-onset^6^ and have lower C-peptide levels^7^, consistent with pancreatic beta cell dysfunction^9^. Earlier studies have reported other rare coding allele(s) in *PAX4* as a cause of monogenic diabetes^8^. However, the high frequency of the variants in the population and a lack of cosegregation with diabetes has led to discussion over whether they are causal for diabetes and if *PAX4* should be included in diagnostic testing panels for monogenic diabetes^10^.

PAX4 is a paired-homeodomain transcription factor that has been shown to act as a transcriptional repressor of insulin and glucagon promoters^11,12^. In mice, *Pax4* is broadly expressed throughout the developing pancreas^12,13^. *Pax4* homozygous knockout mice die three days postpartum from hyperglycemia, caused by a near complete absence of insulin-producing beta cells^13^. Loss of *Pax4* in mice also leads to an increase in the number of alpha cells and an upregulation of the alpha cell gene *Arx*^13,14^. Conversely, *Arx*^-/-^ mice upregulate *Pax4* and have more beta cells, leading to the model of mutual repression of *Pax4* and *Arx* to direct the development of alpha and beta cell lineages, respectively^14^. Similar mutual repression of *pax4* and *arx* has been detected in zebrafish^15^; however, unlike in mice, *Pax4* is not required for beta cell development^15^. In humans, it is currently unknown whether *PAX4* is required for endocrine cell formation. As *PAX4* variants have been reported as a potential cause of monogenic diabetes^8,16,17^ and are associated with altered T2D risk, investigating their role in human endocrine cell formation may improve our understanding of the mechanism(s) underlying the genetic association and clarify the potential role of *PAX4* variants as a cause of monogenic diabetes.

Here we present detailed human *in vivo* and *in vitro* studies on two different *PAX4* coding alleles, the East Asian population enriched p.Arg192His and a novel protein-truncating variant (PTV) p.Tyr186X identified in a Singapore family with early onset diabetes. We generated three independent human induced pluripotent stem cell (hiPSC) models: i) *PAX4*-knockout and *PAX4* variant isogenic SB Ad3.1 cell lines using CRISPR-Cas9 genome editing; ii) donor-derived cells with p.Arg192Arg, p.Arg192His, p.His192His and p.Tyr186X genotypes; and iii) genotype-corrected donor-derived cells, and differentiated all lines into pancreatic beta-like cells (BLCs) using two different protocols. Consistently, we found that *PAX4* deficiency and/or loss-of-function to result in derepression of alpha cell genes, leading to the formation of polyhormonal endocrine cells *in vitro*, with reduced total insulin content. This phenotype was confirmed independently in the human beta cell line EndoC-βH1, and could be reversed in donor-derived hiPSC lines through correction of diabetes-associated *PAX4* alleles. We conclude that, whilst PAX4 is not essential for *in vitro* stem cell differentiation to BLCs, both *PAX4* haploinsufficiency and loss-of-function coding alleles increase the risk of developing diabetes by negatively impacting human beta cell development and insulin secretion. Our observations are consistent with rare *PAX4* alleles resulting in haploinsufficiency being insufficient to cause fully penetrant monogenic diabetes but increasing the risk for T2D.

## Results

### Carriers of the *PAX4* p.Arg192His T2D-risk allele exhibit decreased beta cell function

We recruited a total of 183 non-diabetic individuals and assessed their pancreatic beta cell function by a frequently sampled intravenous glucose tolerance test. Carriers of the T2D-risk allele (p.His192) had a decreased acute insulin response to glucose (AIRg, p=0.002), which remained significant after adjusting for age, sex and BMI (padj=0.04) (Fig. 1a). There were no differences in insulin sensitivity (Si, p=0.105) or disposition index (DI, p=0.203) between the two groups (Fig. 1a). HOMA-B, a measurement of beta cell function, was significantly reduced in p.His192 allele(s) carriers (p=0.005) but was no longer significant after adjusting for age, gender and BMI (padj=0.075). A subset of the recruited individuals [n=57] then underwent an oral glucose tolerance test (OGTT) which revealed higher fasting and 2-hour glucose levels, as well as a lower ratio of area under the curve (AUC) for insulin:glucose (Fig. 1b-d). HOMA-B was significantly poorer during the OGTT in the p.His192 risk allele(s) carriers (132.5±56.2) compared to controls (190.6±98.5) whether unadjusted (p=0.008) or adjusted for age, gender and BMI (padj=0.007). There were no differences in fasting, 2-hour, or AUC glucagon (Extended Data Fig. 1a-c). However, there was a significant decrease in the difference between fasting and 2-hour glucagon (delta glucagon), suggesting carriers of p.His192 have less glucagon suppression (Extended Data Fig. 1d). Consistent with the insulin sensitivity measurement (Fig. 1a), there was no difference in HOMA-IR (Extended Data Fig. 1e). As loss of *Pax4* impacts enteroendocrine cell formation in mice^18^, we also measured GLP-1 in p.His192 carriers and found no significant differences in GLP-1 level (Extended Data Fig. 1f-h). Together, the clinical data are consistent with increased T2D-risk via defects in pancreatic beta cell mass and/or function.

**Figure 1.**
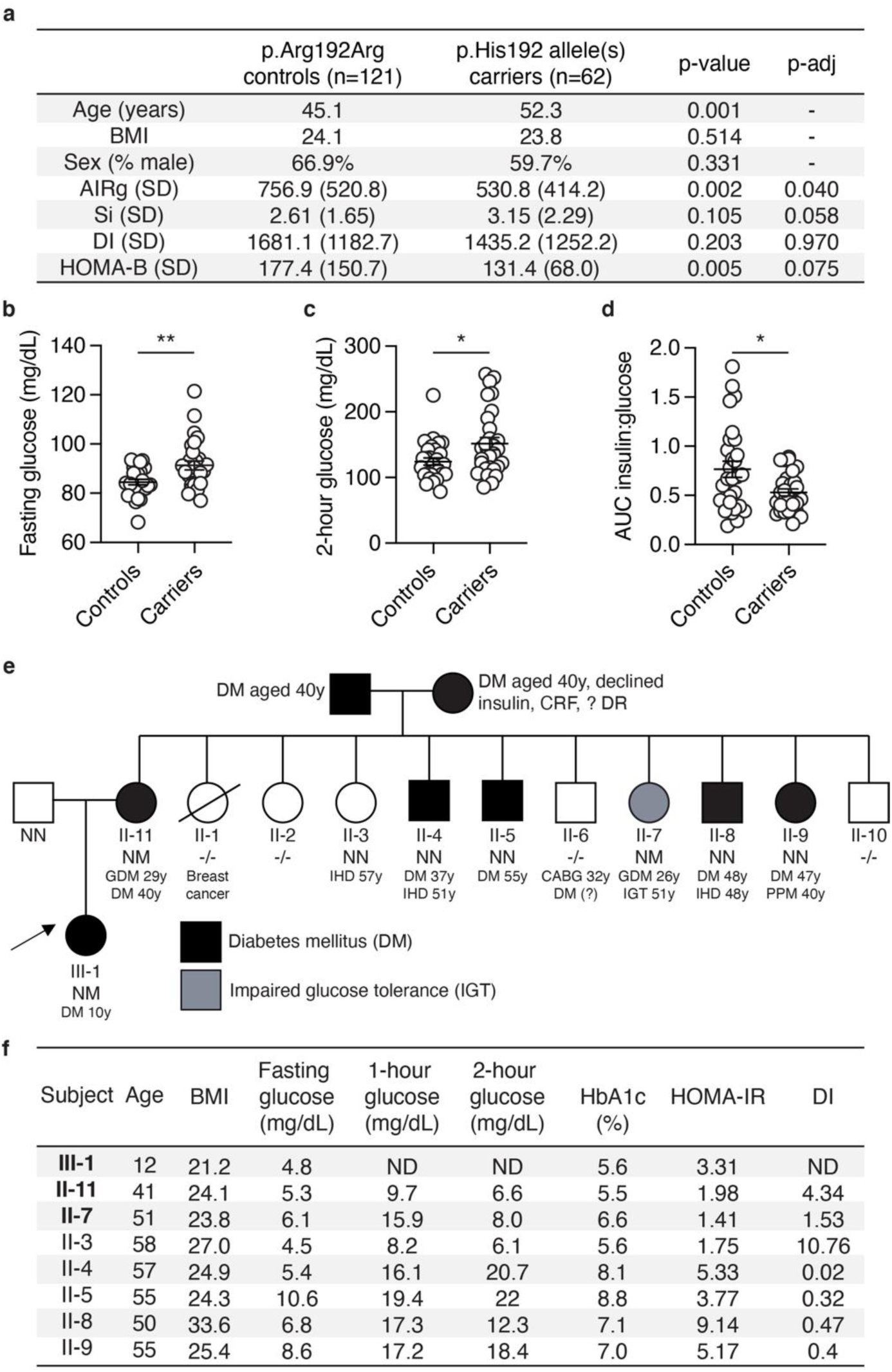
Reduced pancreatic beta cell function in carriers of diabetes-associated *PAX4* variants. (**a**) Mean age and BMI of heterozygous and homozygous carriers of p.His192 *PAX4* allele (n=62) and homozygous p.Arg192Arg controls (n=121) who underwent frequently sampled intravenous glucose tolerance tests to measure Acute Insulin Response to glucose (AIRg), insulin sensitivity (Si), disposition index (DI), and HOMA-B. Unadjusted p-value and adjusted p-value (adjusted for age, sex and BMI) are indicated in the table for AIRg, Si, DI and HOMA-B. (**b-c**) Plasma glucose level (mg/dL) at the (**b**) fasting and (**c**) 2-hour time points during the oral glucose tolerance test (OGTT) of heterozygous or homozygous p.Arg192His carriers (n=29) and p.Arg192Arg controls (n=28). (**d**) Ratio of area under the curve (AUC) insulin to glucose during the 2-hour oral glucose tolerance test. (**e**) Family pedigree of a Singaporean family with a novel p.Tyr186X *PAX4* variant. NN, wild-type; NM, heterozygotes; -/-, genotype not accessible. An arrow indicates the proband (III-1). Age of diagnosis for diabetes mellitus (DM), gestational diabetes mellitus (GDM), ischemic heart disease (IHD), coronary artery bypass grafting (CABG), impaired glucose tolerance (IGT), permanent pace-maker implantation (PPM), chronic renal failure (CRF), and diabetic retinopathy (DR). (**f**) Summary of measures of beta cell function between family members in (**e**) during a 2-hour 75 g glucose oral glucose tolerance test. Carriers of the p.X186 allele are in bold. HbA1c, hemoglobin A1c; HOMA-IR, homeostatic model assessment of insulin resistance (value >2 indicates insulin resistance); DI, disposition index (Matsuda); ND, not done. Data are presented as mean±SEM in Fig. 1b-d. Statistical analyses were performed using unpaired t-test. *p<0.05, **p<0.01.

### Identification of a novel *PAX4* protein truncating variant p.Tyr186X in a family with early onset diabetes

A female proband (III-1) of Singapore Chinese ethnicity was diagnosed with early-onset diabetes at the age of 10 years (random glucose 17 mmol/L) (Fig. 1e), verified to be GAD antibody negative, and had detectable C-peptide (1.2 nmol/L). Upon diagnosis, she was treated with a basal bolus insulin regimen and metformin for two weeks before being switched to metformin-alone treatment. Following lifestyle modifications, she lost weight (from 53.1 kg, BMI 25.3 kg/m^2^ to 49.5 kg, BMI 23.6 kg/m^2^) and nine months post-diagnosis her HbA1c was 7.1% (8.7 mmol/L). The early diabetes-onset, lack of evidence for type 1 diabetes, and persistence of diabetes despite weight loss prompted further assessment in the family (Fig. 1e). II-11 was diagnosed with gestational diabetes (GDM) at the age of 29 years when she was pregnant with the proband. At age 40 years, while being asymptomatic for diabetes, an OGTT confirmed a diagnosis of diabetes with a fasting glucose of 5.6 mmol/L and a 2-hour glucose of 11.4 mmol/L.

Genetic testing for monogenic diabetes with a custom Illumina Nextera rapid capture next-generation sequencing panel on an Illumina Miseq sequencing platform (*HNF4A*, *GCK*, *HNF1A*, *PDX1*, *HNF1B*, *NEUROD1*, *KLF11*, *CEL*, *PAX4, INS, ABCC8*, *KCNJ11*) was performed on members of the family who were recruited for the clinical study. A novel (not reported in gnomAD or ClinVar, date accessed Feb 2022), heterozygous *PAX4* mutation (c.555_557dup) predicted to result in a truncated protein (p.Tyr186X) was identified in the proband (III-1), the mother (II-11) and a female member of the family (II-7) (Fig. 1e). No rare coding variants were detected in the other genes tested.

The other female heterozygous p.Tyr186X variant carrier (II-7; BMI 23.8 kg/m^2^) had a history of GDM (age 26 years) and at the time of study (age 51 years) had impaired glucose tolerance (IGT). Family members (II-4, II-5, II-8 and II-9) with diabetes and another non-diabetic female family member (II-3, BMI 27.0 kg/m^2^) did not carry the variant. Unfortunately, the proband’s grandparents, both diagnosed with diabetes at the age of 40 years, declined to take part in the study.

Given the high prevalence of diabetes in the family and both maternal grandparents having diabetes, we evaluated measures of insulin resistance (HOMA-IR) and beta cell function (DI) in family members with and without the *PAX4* p.Tyr186X variant. Family member II-3, who does not have diabetes and does not carry the p.Tyr186X variant, has the highest beta cell function as measured by the DI whilst those carrying the *PAX4* variant (II-11 and II-7) have markedly reduced function (Fig. 1f). Of note, the family members with diabetes who do not carry the *PAX4* variant all displayed evidence of insulin resistance (HOMA-IR >2) and low DI, consistent with T2D (Fig. 1f). Taken together, these findings are insufficient to provide support for the p.Tyr186X variant as the cause of monogenic diabetes in this family but are consistent with the *PAX4* variant being associated with decreased pancreatic beta cell function.

To further explore the role of rare coding variants in the *PAX4* gene on T2D risk, we accessed aggregated gene-level exome-sequencing association data from 52K individuals deposited in the Common Metabolic Disease Portal (https://t2d.hugeamp.org) and in 281,852 individuals from UKBioBank (https://www.ukbiobank.ac.uk/). Burden and Sequence Kernel Association Test (SKAT) analyses computed using a series of genotype filters and masks provided nominal evidence for a gene level association that is independent of the p.Arg192His variant (Supplementary Table 1a-b).

### Loss of *PAX4* alters hormone gene regulation, reduces insulin secretion function and total insulin content in a human beta cell model

To evaluate the consequence of *PAX4* loss on beta cell function, we first performed siRNA- and shRNA-mediated knockdown of *PAX4* in human EndoC-βH1 cells (Fig. 2a). Transient knockdown of *PAX4* using siRNAs significantly reduced *PAX4* transcript expression (Fig. 2b) and glucose-stimulated insulin secretion (GSIS) in EndoC-βH1 cells (Fig. 2c). To model a chronic loss of *PAX4* expression, we generated stable knockdown *PAX4* EndoC-βH1 cells via lentiviral transduction of shRNA followed by antibiotic selection (Fig. 2a). sh*PAX4* EndoC-βH1 cells had reduced *PAX4* transcript level compared to shScramble control cells (Fig. 2d). Long-term knockdown of *PAX4* completely abolished GSIS compared to shScramble control cells (Fig. 2e), accompanied by reduced total insulin content in sh*PAX4* EndoC-βH1 cells (Fig. 2f). While there was no difference in *INS* transcript (Fig. 2g), loss of *PAX4* increased *GCG* transcript by 6.2-fold in sh*PAX4* EndoC-βH1 cells (Fig. 2h), consistent with PAX4 being a repressor of *GCG* expression^12,19^.

**Figure 2.**
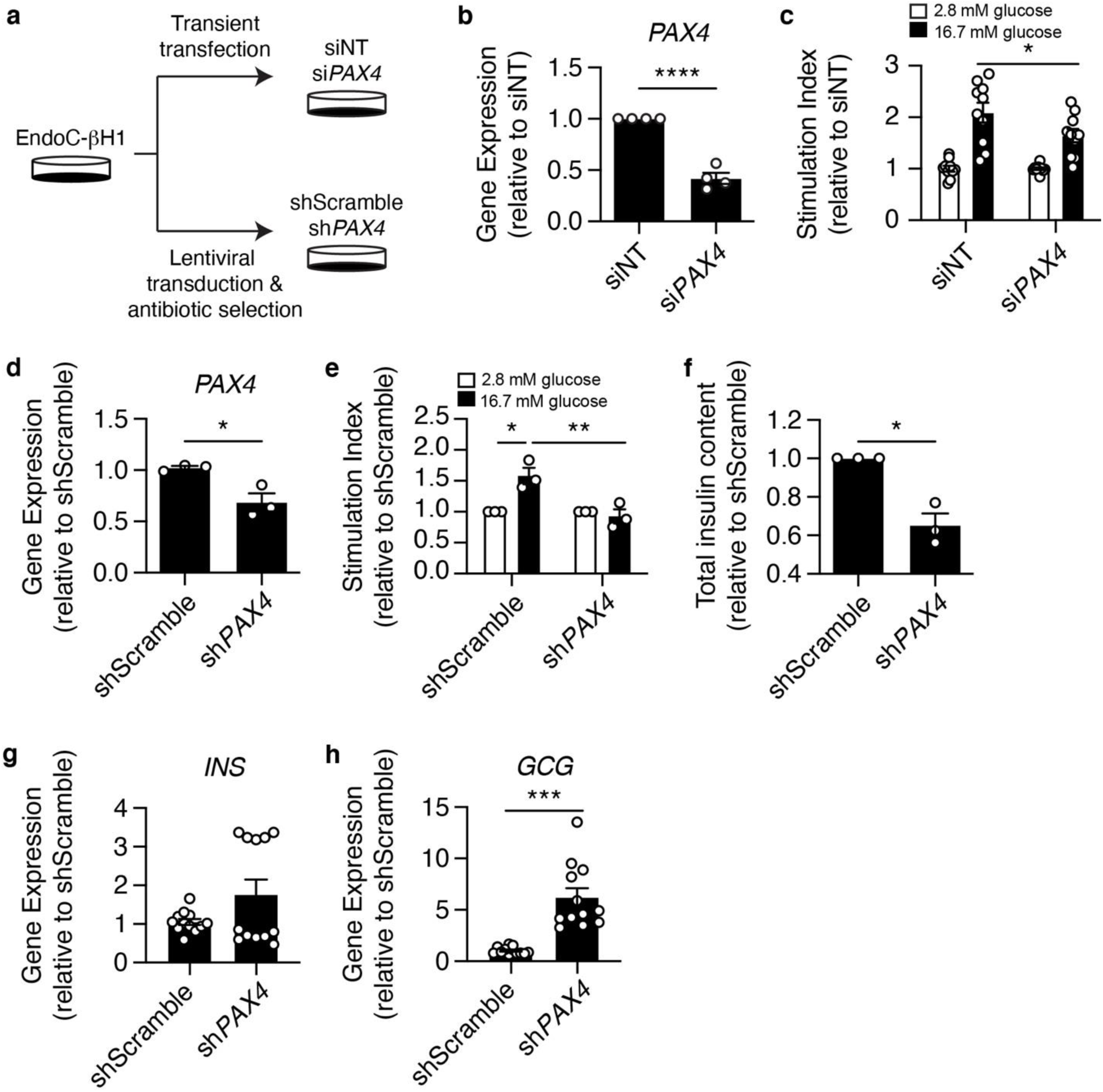
*PAX4* knockdown and knockout impairs glucose-stimulated insulin-secretion and reduces insulin content in human EndoC-βH1 cells. (**a**) Experimental design for *PAX4* knockdown approaches using siRNA and shRNA in EndoC-βH1 cells. (**b**) *PAX4* gene expression following transient transfection of si*PAX4* and non-targeting (siNT) control in EndoC-βH1 cells. (**c**) Glucose-stimulated insulin secretion of 2.8 mM and 16.7 mM glucose in siNT and si*PAX4* EndoC-βH1 cells, normalized to total protein then to 2.8 mM glucose. (**d**) *PAX4* gene expression in *PAX4*-knockdown (sh*PAX4*) and control (shScramble) EndoC-βH1 cells following six passages of antibiotic selection. (**e**) Glucose-stimulated insulin secretion assay comparing sh*PAX4* and shScramble EndoC-βH1 cells, normalized to total DNA and then to 2.8 mM glucose. (**f**) Relative fold change in total insulin content in sh*PAX4* and shScramble EndoC-βH1 cells, normalized to total DNA content. (**g**) *INS* transcript expression in shScramble and sh*PAX4* EndoC-βH1 cells. (**h**) *GCG* transcript expression in shScramble and sh*PAX4* EndoC-βH1 cells. Data are presented as mean±SEM. Statistical analysis of two samples was performed by paired t test or a two-way ANOVA for comparison of multiple groups. *p<0.05, ***p<0.001, ****p<0.0001. n=3-12.

### *PAX4* knockout in hiPSC-derived BLCs causes derepression of alpha cell gene expression

While *PAX4* transcript and protein can be detected in rat^20^ and human islets^21^, its expression is most abundant during embryonic development^12,13^, suggesting that *PAX4* variants may mediate disease risk early on during embryonic development. Homozygous *Pax4* knockout mice die within three days of birth and have a near complete loss of pancreatic beta cells^13^. Whether *PAX4* is similarly required for the formation of human beta cells is unknown. We generated *PAX4* homozygous null isogenic hiPSC lines (*PAX4*^+/+^; *PAX4*^-/-^) using CRISPR-Cas9 (Fig. 3a) and two single guide RNAs (sgRNAs) designed for exons 2 and 5 (encoding the paired-domain and homeodomain) of the *PAX4* gene (Extended Data Fig. 2a). Two independent cell lines had a homozygous deletion for amino acids 64 through 200, whilst the other cell line was compound heterozygous for two premature stop codons at amino acids 61 and 74, respectively (data not shown). Three independent, unedited hiPSC lines generated during the CRISPR-Cas9 process provided control *PAX4*^+/+^ lines (Fig. 3a). All six hiPSC lines were differentiated towards BLCs using a seven-stage protocol^22^ (Fig. 3b; Protocol A). Flow cytometry analysis of CXCR4+ and SOX17+ definitive endoderm (DE) cells determined that there was no defect in the formation of DE (Extended Data Fig. 2b-c). As *PAX4* is a transcription factor, RNA-seq analysis was used to determine the transcriptional consequence of *PAX4* knockout. RNA-seq samples (n=8 per genotype) were collected at DE (before *PAX4* expression), pancreatic endoderm (PE), endocrine progenitor (EP) (at the peak of *PAX4* expression) and BLC (Fig. 3c and Supplementary Table 2). The *PAX4* transcript was significantly reduced in *PAX4*^-/-^ PE (padj=4.25E-05), EP (padj=6.15E-05) and BLC (padj=1.27E-06) (Fig. 3c), and the remaining transcripts were missing exons 2 through 5 (Extended Data Fig. 2d).

**Figure 3.**
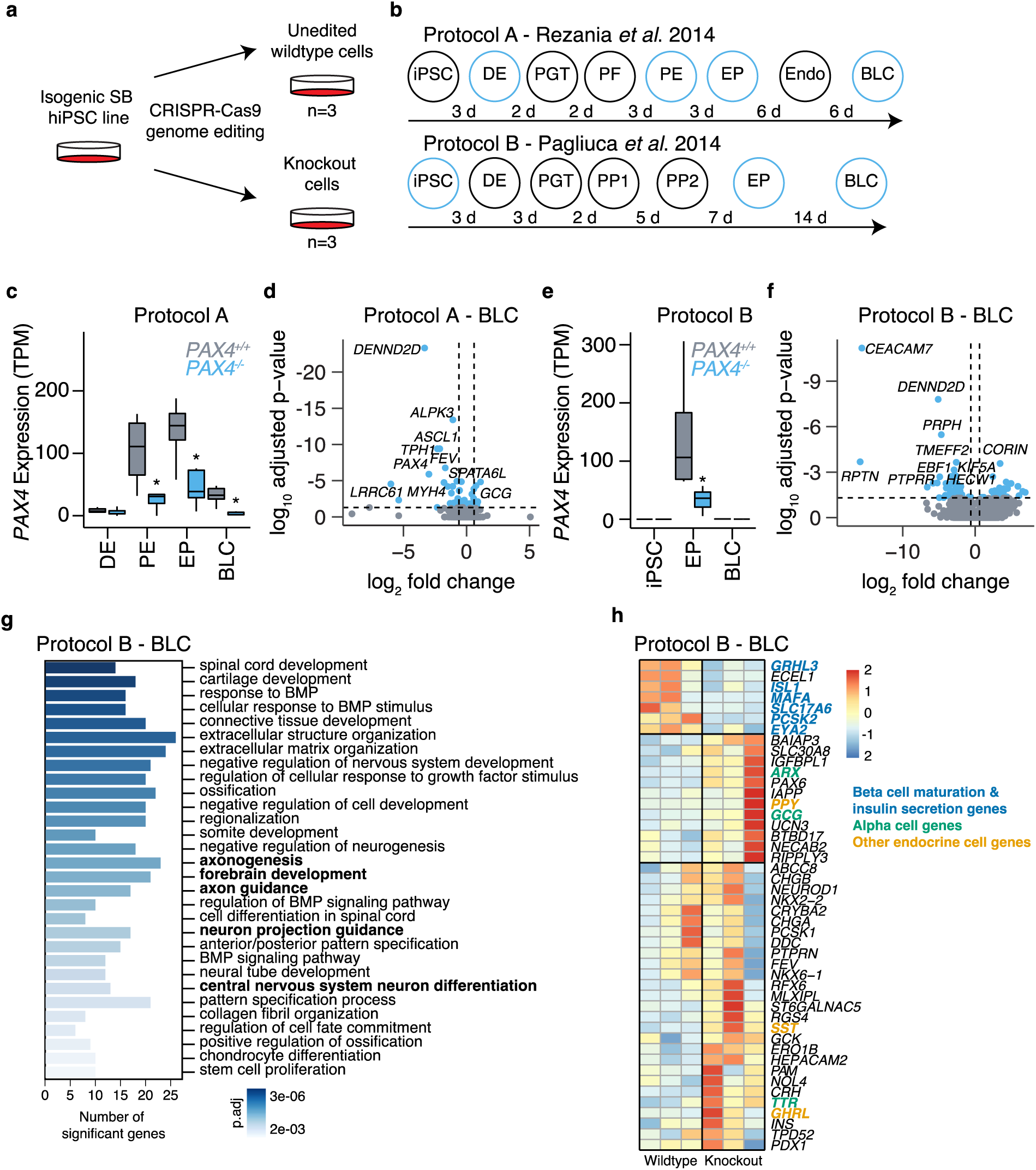
Isogenic *PAX4* knockout human induced pluripotent stem cell lines have defects in endocrine cell formation during *in vitro* differentiation. (**a**) CRISPR-Cas9 genome editing was used to generate three independent isogenic hiPSC *PAX4* homozygous knockout cell lines and three unedited wildtype control cell lines. (**b**) Schematic outline of seven-stage protocol A and six-stage protocol B from human induced pluripotent stem cells (hiPSC), definitive endoderm (DE), primitive gut tube (PGT), posterior foregut (PF) or pancreatic progenitor 1 (PP1), pancreatic endoderm (PE) or pancreatic progenitor 2 (PP2), endocrine progenitor (EP), endocrine (Endo), towards beta-like cells (BLC). RNA-seq samples were collected at the end of stages highlighted in blue. (**c**) Expression of *PAX4* in transcripts per million (TPM) in *PAX4*^+/+^ and *PAX4*^-/-^ cells at DE, PE, EP and BLC derived from Protocol A. (**d**) Volcano plot of differentially expressed genes in *PAX4*^-/-^ versus *PAX4*^+/+^ BLCs derived from Protocol A. The top ten differentially expressed genes are highlighted. (**e**) Expression of *PAX4* in TPM in *PAX4*^+/+^ and *PAX4*^-/-^ cells at hiPSCs, EPs and BLCs derived from Protocol B. (**f**) Volcano plot of differentially expressed genes in *PAX4*^-/-^ versus *PAX4*^+/+^ BLCs derived from Protocol B. The top ten differentially expressed genes are highlighted. (**g**) Gene Ontology (GO) analysis of differentially expressed genes in *PAX4*^-/-^ compared to *PAX4*^+/+^ BLCs from Protocol B. (**h**) Heatmap of relative gene expression for pancreatic endocrine genes in BLCs derived from Protocol B.

Differential expression analysis using DESeq2 showed that the loss of *PAX4* resulted in a de-repression of an alpha cell gene signature (*ARX, GCG, TTR*) and a repression of the endocrine progenitor marker *FEV*^23,24^ in BLCs (Fig. 3d and Extended Data Fig. 2e-g). To confirm our results, we differentiated the same *PAX4*^+/+^ and *PAX4*^-/-^ hiPSC lines into BLCs using a second protocol (Fig. 3b; Protocol B)^25^. Using Protocol B, we found a significant reduction in *PAX4* transcript at the EP stage (Fig. 3e) and a larger number of differentially expressed genes (Fig. 3f and Supplementary Table 3). Gene ontology (GO) biological process analysis of the differentially expressed genes in *PAX4*^-/-^ BLCs revealed a number of GO terms that included the alpha cell gene *ARX*.

Using a curated list of genes involved in beta and alpha cell lineages^26^, we observed directionally consistent, albeit nonsignificant, derepression of alpha cell genes (*ARX*, *GCG*, *TTR*) (Fig. 3h). The expression of delta cell gene (*SST*), epsilon cell gene (*GHRL*) and PP cell gene (*PPY*) was also derepressed in BLCs derived from *PAX4^-/-^* lines (Fig. 3h). Some genes that are involved in beta cell maturation and hormone secretion (*MAFA, ISL1, GRHL3, SLC17A6, PCSK2, EYA2*) were downregulated with the loss of *PAX4* (Fig. 3h). Importantly, *PAX4^+/+^* and *PAX4^-/-^* lines differentiating into BLCs repress pluripotency genes and activate genes involved in endocrine cell fate in a similar manner (Extended Data Fig. 3), suggesting that, unlike in mouse, *PAX4* is not required for human beta cell differentiation *in vitro*. Rather, *PAX4* loss-of-function results in derepression of alpha cell genes and a dysregulation of key endocrine maturation genes in hiPSC-derived BLCs.

### Donor-derived hiPSCs from carriers of the *PAX4* p.Arg192His and p.Tyr186X alleles have defects in endocrine cell differentiation *in vitro*

Having established the effect of *PAX4* loss during *in vitro* beta cell differentiation, we next generated donor-derived hiPSCs from *PAX4* variant carriers and differentiated them into BLCs. Skin biopsies and/or blood samples from recruited non-diabetic donors were used to derive hiPSCs of the following genotypes: homozygous for the *PAX4* p.Arg192 and p.Tyr186 alleles (wildtype), heterozygous for either the p.Arg192His or p.Tyr186X alleles, and homozygous for the p.His192 allele (p.His192His) (Fig. 4a). To account for possible line-to-line heterogeneity in hiPSC-based studies^27^, three independent hiPSC lines were generated from two donors for wildtype cells, four lines from two donors for p.Arg192His, five lines from two donors for p.His192His, and three lines from one donor (II-7; Fig. 1e) for the p.Tyr186X variant (Fig. 4a). All hiPSC lines were characterized via pluripotency immunostaining, teratoma assay, and karyotyping and genotypes were confirmed by Sanger sequencing (data not shown).

**Figure 4.**
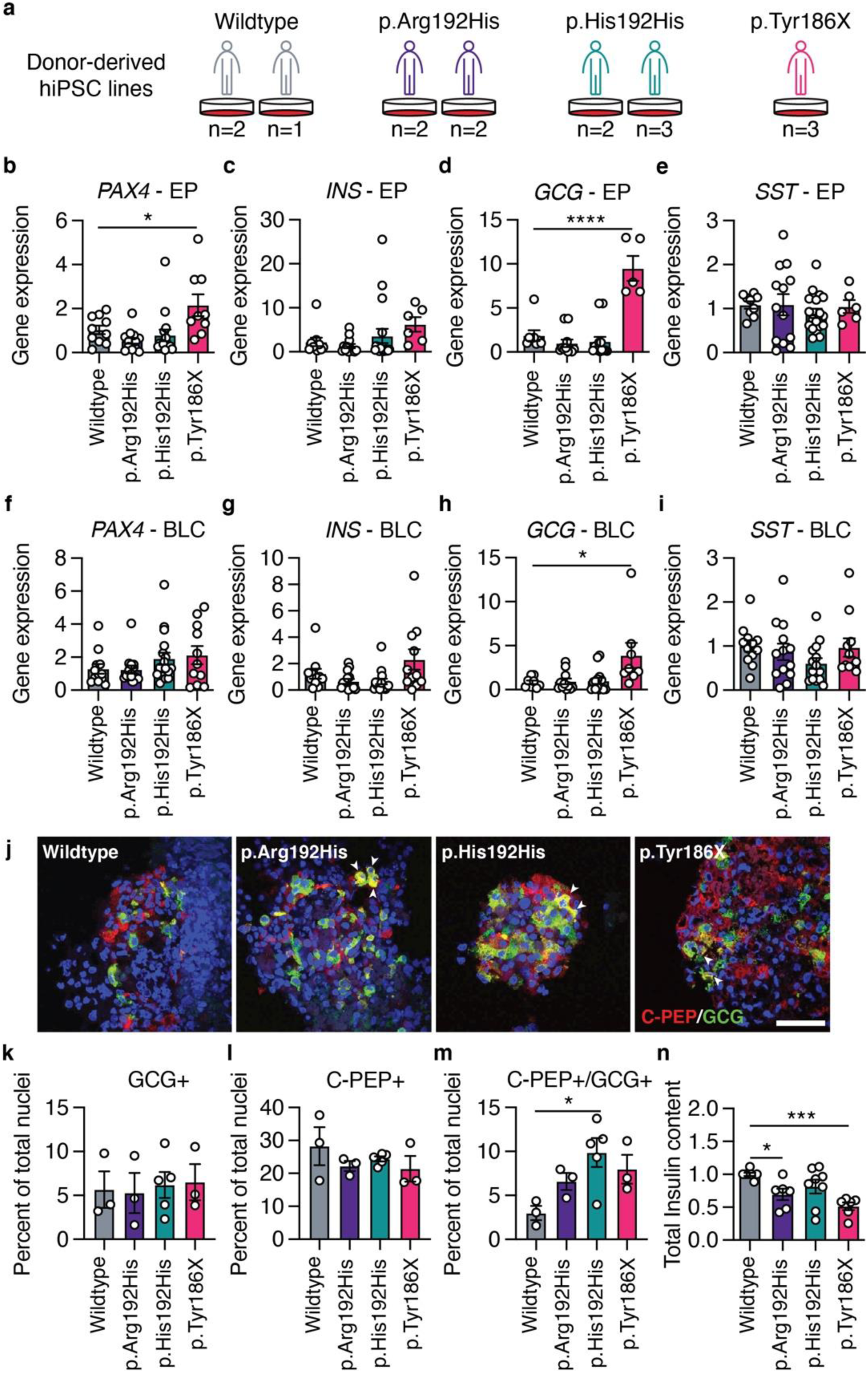
*PAX4* p.Arg192His and p.Tyr186X donor-derived hiPSCs have perturbations in differentiation towards BLCs. (**a**) Donor-derived hiPSC lines were generated from the following genotypes: three lines from two wildtype donors; four lines from two p.Arg192His donors; five lines from two p.His192His donors; and three lines from one p.Tyr186X donor. (**b-e**) Transcript expression of (**b**) *PAX4*, (**c**) *INS*, (**d**) *GCG* and (**e**) *SST* in hiPSC-derived endocrine progenitor (EP) cells using differentiation Protocol B. (**f-i**) Transcript expression of (**f**) *PAX4*, (**g**) *INS*, (**h**) *GCG* and (**i**) *SST* in hiPSC-derived beta-like cells (BLCs) using differentiation Protocol B. (**j**) Representative immunofluorescence images of hiPSC-derived beta-like cells with C-peptide in red, glucagon in green, and nuclei in blue. Arrows indicate C-PEP+/GCG+ double-positive cells. Scale bar: 50 µm. (**k-m**) Quantification of immunofluorescence images for the percentage of cells expressing (**k**) GCG (monohormonal), (**l**) C-PEP (monohormonal) or (**m**) C-PEP+/GCG+ (polyhormonal). (**n**) Total insulin content normalized to total DNA from hiPSC-derived BLCs. Data are presented as mean±SEM. n>3. Statistical analyses were performed using one-way ANOVA. *p<0.05, ***p<0.001, and ****p<0.0001.

We simultaneously differentiated all 15 hiPSC lines into pancreatic BLCs using Protocol B and performed qPCR, flow cytometry and immunostaining analyses. *PAX4* transcript expression was unchanged in carriers of the *PAX4* p.His192 allele but elevated in EPs derived from the p.Tyr186X carrier (Fig. 4b), consistent with transcriptional compensation for the PTV. Heterozygous and homozygous carriers of the *PAX4* p.His192 allele had no measurable differences in *INS*, *GCG*, or *SST* gene expression at the EP or BLC stages (Fig. 4b-i). Similar to the *PAX4*^-/-^ knockout hiPSCs (Fig. 2h), the *PAX4* p.Tyr186X hiPSC lines exhibited a derepression of the *GCG* gene in both EPs and BLCs (Fig. 4d and h), suggesting that the PTV is loss-of-function. Immunostaining of endocrine hormones found no significant differences in GCG+ or INS+ cells (Fig. 4j-l), but there was a significant increase in the number of polyhormonal (C-PEP+/GCG+) BLCs from *PAX4* variant hiPSC lines (Fig. 4j and m). BLCs from the *PAX4* p.Arg192His and p.Tyr186X hiPSCs had lowered total insulin content (Fig. 4n), which is consistent with the *in vivo* clinical data from *PAX4* variant carriers (Fig. 1a and f). Together, these data suggest that both *PAX4* alleles result in a loss-of-function due to reduced *PAX4* gene dosage and/or altered PAX4 transcriptional activity, negatively affecting endocrine cell differentiation.

### *PAX4* p.Arg192His and p.Tyr186X alleles reduce the expression and/or function of PAX4 protein

The PAX4 protein consists of two functional domains, paired and homeodomain, that are responsible for DNA binding, and two nuclear localization sequences (NLS) (Fig. 5a)^28,29^. Both p.Arg192His and p.Tyr186X variants are located within the functional homeodomain of the PAX4 protein (Fig. 5a). As the crystal structure of PAX4 protein has not been elucidated, we obtained the predicted three-dimensional molecular arrangement of PAX4 protein (AF-O43316-F1-model_v2) from the AlphaFold database^30,31^. The p.Arg192 residue is located within a hydrophobic pocket (Extended Data Fig. 4a), suggesting that substitution to an uncharged histidine may alter the DNA-binding function of the PAX4 protein. The *PAX4* p.Tyr186X (c.557-559 GTA duplication) variant causes a frameshift, leading to the introduction of a premature stop codon at amino acid position 186 (Fig. 5a and Extended Data Fig. 4a). Transcripts containing PTVs, such as *PAX4* p.Tyr186X, may undergo nonsense mediated decay (NMD), resulting in haploinsufficiency^32^. To test whether the *PAX4* variants undergo NMD, we performed allelic-specific qPCR following treatment with the NMD inhibitor cycloheximide (CHX)^33,34^. Treatment of hiPSC-derived BLCs with CHX overnight stabilized the p.X186 allele (Fig. 5b) but had no effect on p.His192 (Fig. 5c), confirming NMD of p.X186 and supporting *PAX4* haploinsufficiency as the mechanism for the p.Tyr186X variant.

**Figure 5.**
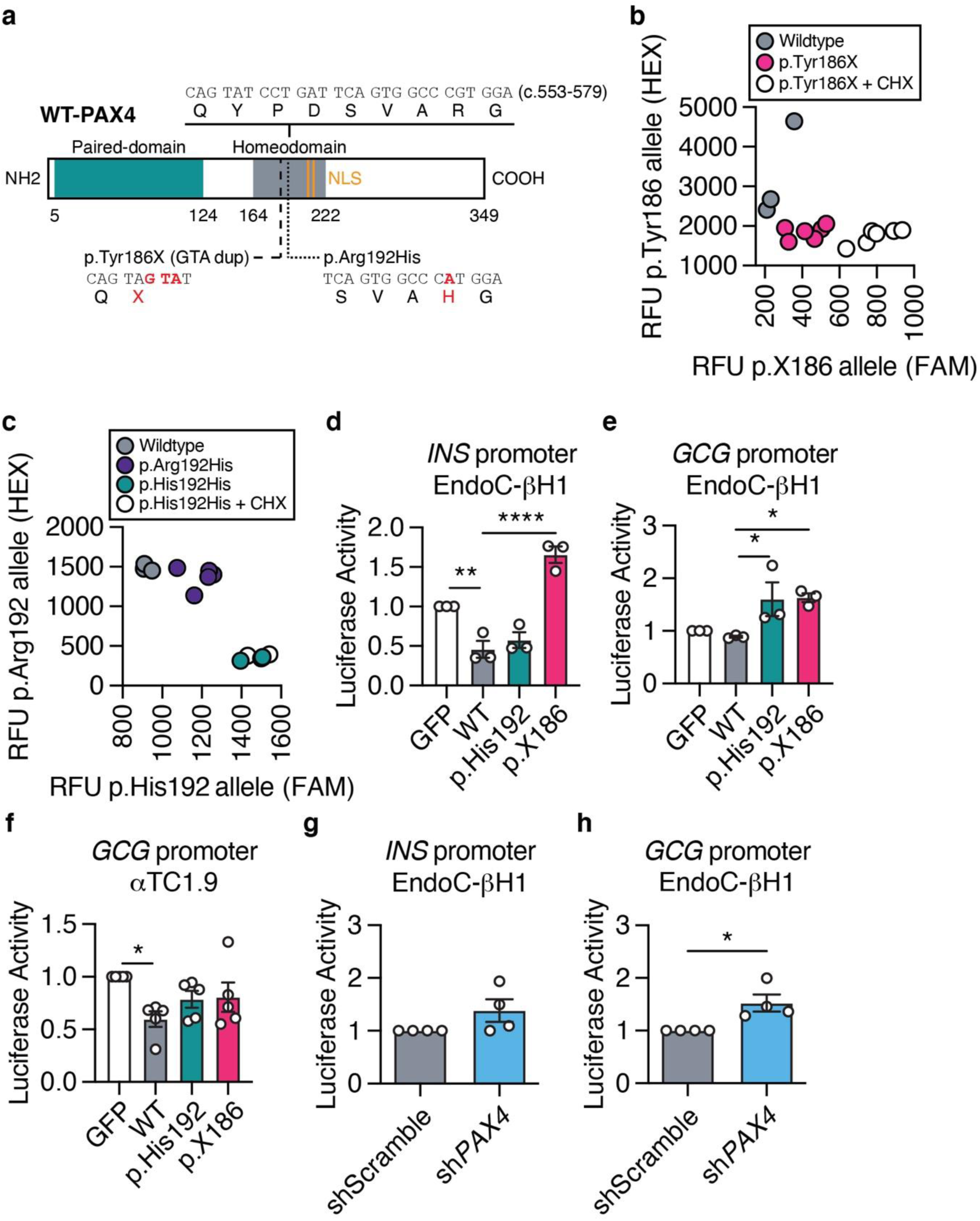
Molecular function of PAX4 variants in human EndoC-βH1 cells. (**a**) Illustration of human full-length wildtype (WT) PAX4 protein. Functional domains are depicted in green (paired-domain) and grey (homeodomain). Nuclear localization sequences (NLS; orange) are located at amino acids 206-210 and 212-216. Part of the homeodomain sequence that contains the *PAX4* variants (c.553-579) and the amino acid changes of p.Tyr186X and p.Arg192His are highlighted. (**b)** Allele-specific qPCR of *PAX4* transcript for p.Tyr186 and p.X186 alleles following cycloheximide (CHX) treatment. (**c**) Allele-specific qPCR of *PAX4* transcript for p.Arg192 and p.His192 alleles following CHX treatment. (**d-e**) Luciferase activity of (**d**) *INS* and (**e**) *GCG* gene promoters in EndoC-βH1 cells overexpressing WT-PAX4, p.His192 and p.X186. (**f**) Luciferase activity of the *GCG* gene promoter in αTC1.9 cells overexpressing WT-PAX4, p.His192 and p.X186. (**g-h**) Luciferase activity of (**g**) *INS* and (**h**) *GCG* gene promoters in shScramble and sh*PAX4* EndoC-βH1 cells. Data are presented as mean±SEM. Statistical analyses were performed by t-test or two-way ANOVA. n>3. *p<0.05, **p<0.01, ****p<0.0001.

To understand the consequence of *PAX4* variants on protein function, we performed a series of *in vitro* assays using overexpression of tagged WT and mutant (p.His192 and p.X186) PAX4 protein (Extended Data Fig 4b). Western blot analyses demonstrated successful PAX4 overexpression detected by PAX4 antibody^35^ and V5 tag expression (Extended Data Fig. 4c). Overexpression of the *PAX4* variants in AD293 cells confirmed that the p.His192 allele does not prevent nuclear localization and that any p.X186 protein that escaped NMD remained trapped in the cytoplasm due to the loss of downstream NLS (Extended Data Fig. 4d). We observed fewer PAX4 antibody-positive cells in p.X186 transfected AD293 cells, despite no difference in overall transfection efficiency (% GFP+), consistent with decreased stability of any truncated protein produced by the PTV (Extended Data Fig. 4e). Treating AD293 cells overexpressing the *PAX4* constructs with the proteasomal inhibitor MG132 revealed an accumulation of the p.X186 protein compared to wildtype or p.His192, demonstrating that the overexpressed truncated protein is subject to proteasomal degradation (Extended Data Fig. 4f-g).

It was previously reported that *PAX4* p.His192 results in defective transcriptional repression of human *INS* and *GCG* gene promoters^12^. In EndoC-βH1 cells, overexpression of both WT and p.His192 PAX4 proteins resulted in a significant repression of *INS* promoter activity (Fig. 5d). Although the p.X186 variant most likely results in NMD and haploinsufficiency, any translated protein was unable to repress *INS* promoter activity (Fig. 5d). WT PAX4 protein did not repress the *GCG* gene promoter in EndoC-βH1 cells (Fig. 5e) but did so in the rodent alpha cell model αTC1.9 (Fig. 5f), consistent with cell-type specific regulation of gene expression by PAX4. Both *PAX4* p.His192 and p.X186 resulted in a derepression of the *GCG* promoter in beta cells (Fig. 5e) and a loss of repression activity in alpha cells (Fig. 5f). Luciferase assays for *INS* gene promoter activity were unchanged in EndoC-βH1 cells following *PAX4* (sh*PAX4*) knockdown (Fig. 5g). However, sh*PAX4* EndoC-βH1 cells had significantly increased *GCG* promoter activity (Fig. 5h), consistent with the loss-of-repression of *GCG* promoter activity observed in the presence of p.His192 or p.X186. Taken together, our studies demonstrate that *PAX4* p.Arg192His and p.Tyr186X variant proteins have altered expression and/or transcriptional activity.

### hiPSC-derived EPs have a distinct metabolic gene signature and exhibit a bioenergetics switch from glycolysis to oxidative phosphorylation

To evaluate the overall impact of diabetes-associated *PAX4* gene variants on the global transcriptome of human pancreatic cells, RNA-seq was performed on *PAX4* variant carrier donor-derived hiPSCs across four differentiation time points using Protocol B: hiPSCs, PP2 cells, EPs and BLCs (Supplementary Table 4). Uniform Manifold Approximation and Projection (UMAP) analyses of a total of 153 RNA samples demonstrated that samples were clustered based on differentiation day (i.e., largest source of variation is developmental time point) (Fig. 6a), indicating that the differentiation protocol is robust in directing the hiPSCs toward BLCs. Volcano plots comparing the *PAX4* p.Arg192His, p.His192His or p.Tyr186X against wildtype *PAX4* donor-derived hiPSCs demonstrated that most differentially expressed genes were upregulated at the EP stage (Fig. 6b), coinciding with the peak of *PAX4* expression (Fig. 3c and e). Principal Component Analysis (PCA) revealed that EPs derived from *PAX4* variants clustered more closely to each other than to wildtype *PAX4* (Fig. 6c), suggesting that the two *PAX4* variants shared transcriptional similarity. Gene enrichment analyses of relevant biological processes of the differentially expressed genes in the *PAX4* p.His192His and p.Tyr186X EPs revealed an association with metabolic processes and cellular response to stress (Fig. 6d). Between the *PAX4* p.His192His and p.Tyr186X genotypes, there were 2012 and 452 genes in common within the “metabolic processes” and “cellular response to stress”, respectively, most of which were elevated in expression in the *PAX4* variant lines (Extended Data Fig. 5a-b).

**Figure 6.**
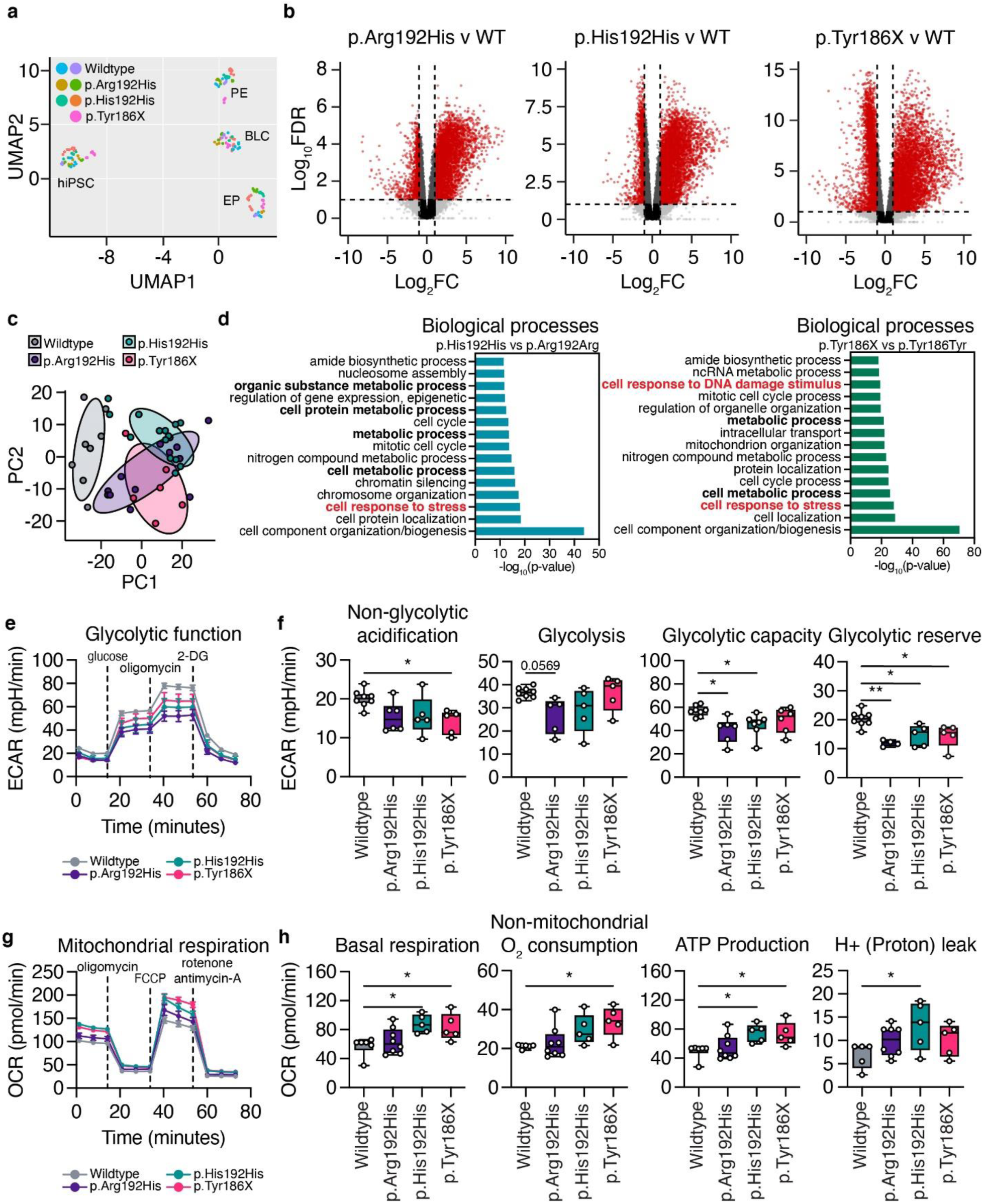
Metabolic seahorse assays revealed alterations in glycolysis function and oxidative phosphorylation in the presence of the *PAX4* p.Arg192His or p.Tyr186X variants. (**a**) Uniform Manifold Approximation and Projection (UMAP) of 153 RNA samples at the hiPSC, PE, EP and BLC stages of *in vitro* differentiation using Protocol B. (**b**) Volcano plots [Log_10_FDR and log_2_(FC)] demonstrating pairwise comparisons of p.Arg192His, p.His192His, and p.Tyr186X against wildtype, respectively. Red circles represent transcripts with log2FC <-2 or >2 and p<0.05. (**c**) Principal component analysis (PCA) of RNA-seq data for *PAX4* donor hiPSC-derived EP cells. PC1: 35%; PC2: 11%. (**d**) Gene ontology (GO) analysis of differentially expressed genes in EP comparing p.His192His against p.Arg192Arg (*PAX4*^+/+^) or p.Tyr186X against p.Tyr186Tyr (*PAX4*^+/+^), FC<0.67 or FC>1.5. The bars denote –log10(p-value) with FDR<0.05. (**e**) Extracellular acidification rate (ECAR) measurements of wildtype, p.Arg192His, p.His192His, and p.Tyr186X EP cells following a sequential addition of glucose, oligomycin, and 2- deoxyglucose (2-DG). (**f**) Non-glycolytic acidification, glycolysis, glycolytic capacity, and glycolytic reserve measurements during the ECAR of wildtype, p.Arg192His, p.His192His, and p.Tyr186X EP cells. (**g**) Oxygen consumption rate (OCR) measurements of wildtype, p.Arg192His, p.His192His, and p.Tyr186X EP cells following a sequential addition of oligomycin, FCCP, rotenone and antimycin-A. (**h**) Basal respiration, non-mitochondrial O_2_ respiration, ATP production, and H+ (proton) leak of wildtype, p.Arg192His, p.His192His, and p.Tyr186X EP cells. Data are presented as mean±SEM. n>3. Statistical analyses were performed by one-way ANOVA. *p<0.05, **p<0.01.

To further assess a potential defect in metabolism, we performed a Seahorse XFe96 Glycolysis Stress Test. The glycolysis stress test showed that EPs carrying one or two p.His192 risk alleles had lower glycolytic function, including glycolytic capacity and glycolytic reserve, and a modest downregulation of glycolysis (Fig. 6e-f). In addition, EPs carrying the p.Tyr186X risk allele had decreased glycolytic reserve and non-glycolytic acidification (Fig. 6f). Next, we hypothesized that EPs would seek alternative metabolic processes to compensate for the reduction in energy production, such as oxidative phosphorylation through mitochondrial respiration. Mitochondrial function was measured via oxygen consumption in EPs using the Seahorse XFe96 analyzer. There was an increase in oxidative phosphorylation activity in EPs harboring *PAX4* variants (Fig. 6g), including basal respiration, non-mitochondrial O_2_ consumption, ATP production, and H^+^ (proton) leak (Fig. 6h). Overall, EPs carrying *PAX4* diabetes risk alleles demonstrated a bioenergetic switch from glycolysis to oxidative phosphorylation.

To investigate if the altered metabolic gene expression and bioenergetics profile contributed to beta cell maturation from the EP stage, we treated differentiating cells with the antioxidant N-acetylcysteine (NAC)^36^ from EP (when *PAX4* expression peaks) to BLC stage and extracted total insulin for assessment. *PAX4* p.His192His and p.Tyr186X carrying BLCs revealed only a modest upregulation in total insulin content (Extended Data Fig. 6a-b), suggesting that the alleviation of oxidative stress is insufficient to rescue the total insulin content in BLCs. We postulate that the metabolic signature observed in our donor-derived hiPSC model reflects the physiological status of the EPs rather than being the immediate cause for the dysregulation of beta cell development and maturation.

### Correction of *PAX4* risk alleles in donor-derived hiPSCs with CRISPR-Cas9 rescues dysregulated endocrine gene expression and metabolic phenotypes

Next, we used CRISPR-Cas9 to correct the donor-derived hiPSC lines and to generate *PAX4* variant isogenic hiPSC lines. We designed sgRNA#3 to target the donor-derived homozygous p.His192His line and provided the homology-directed repair (HDR) template to correct the rs2233580 T2D-risk allele (Fig. 7a). The II-11 donor-derived hiPSC line that is heterozygous for a GTA duplication was corrected with sgRNA#4 and an HDR template (Fig. 7b). From the CRISPR-Cas9 genome editing pipeline, we generated two corrected p.Arg192Arg non-risk and two uncorrected p.His192His hiPSC lines (Fig. 7c). From the II-11 donor-derived line, four corrected p.Tyr186Tyr and four uncorrected p.Tyr186X hiPSC lines were derived (Fig. 7c). All the corrected and uncorrected lines were differentiated towards BLCs using Protocol B, followed by RNA-seq analyses and assessment of total insulin content (Fig. 7c).

**Figure 7.**
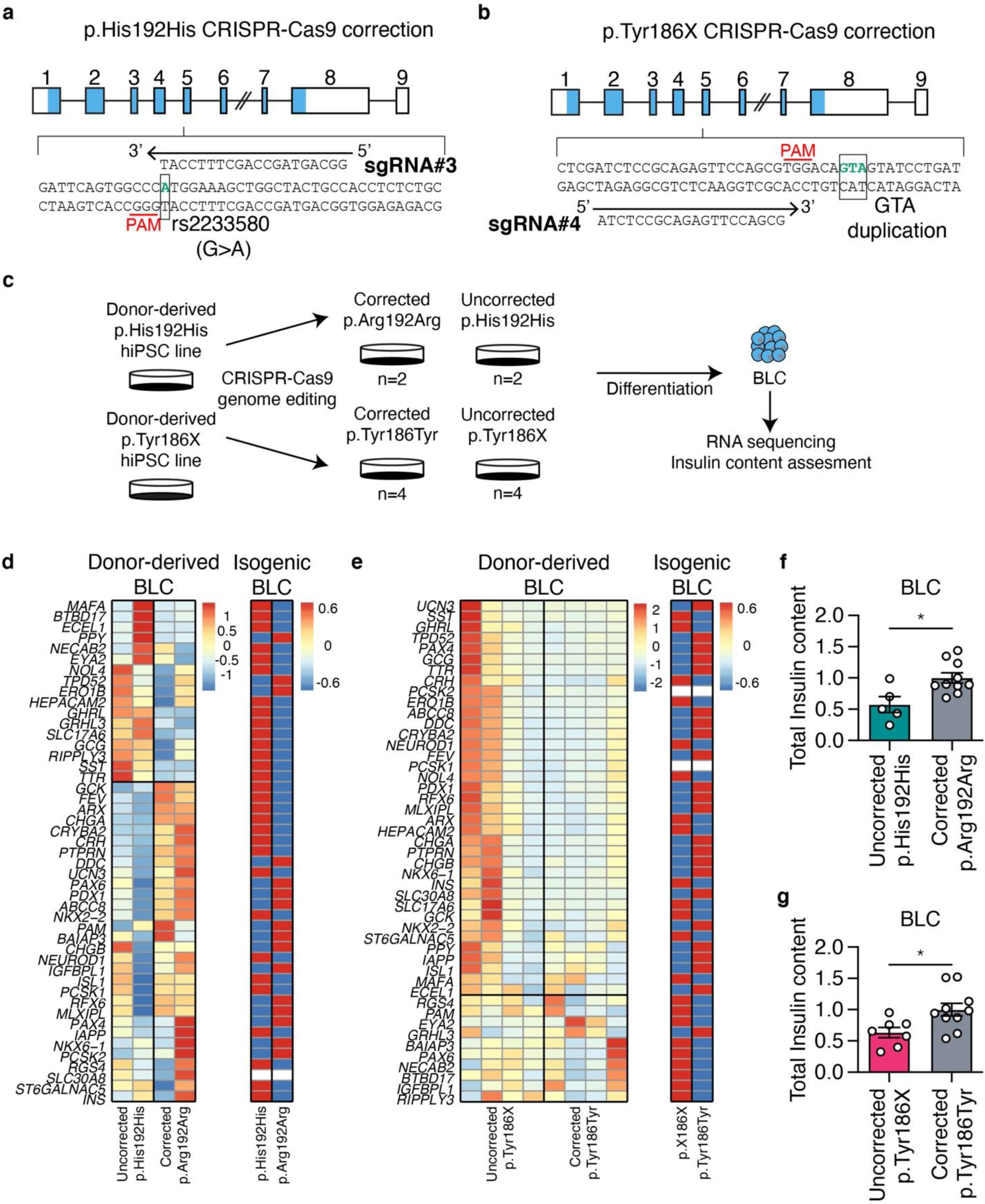
CRISPR-correction of p.Arg192His or p.Tyr186X allele demonstrated rescue in beta cell identity and total insulin content. (**a**) CRISPR-Cas9 gene correction strategy for p.His192His genotype. (**b**) CRISPR-Cas9 gene correction strategy for p.Tyr186X genotype. (**c**) Experimental design for CRISPR-Cas9 mediated gene correction and differentiation strategies. (**d-e**) Targeted heatmap of selected beta and alpha specific genes comparing uncorrected and corrected donor-derived BLCs and isogenic BLCs for (**d**) p.His192 and (**e**) p.X186 alleles. (**f-g**) Total insulin content of BLCs derived from (**f**) uncorrected p.His192His and corrected p.Arg192Arg and (**g**) uncorrected p.Tyr186X and corrected p.Tyr186Tyr. n>3. Data are presented as mean±SEM. Statistical analyses were performed by Student’s t-test, *p<0.05.

Consistent with the BLCs derived from *PAX4*^-/-^ hiPSCs (Fig. 3), homozygous p.His192His donor-derived BLCs had higher expression of non-beta cell genes *GCG*, *TTR*, *SST,* and *PPY* (Fig. 7d and Supplementary Table 5). Correction of *PAX4* p.His192 variant to p.Arg192 rescued the expression of several beta cell genes, such as *RFX6*^37^, *ABCC8*^38^ and *SLC30A8*^39^ (Fig. 7d), which are involved in insulin content and secretion. Isogenic hiPSC-derived BLCs homozygous for the p.His192 allele had directionally consistent changes in gene expression (Fig. 7d), confirming that p.His192 is the cause of perturbed beta cell gene expression. The differences between BLCs derived from corrected or uncorrected *PAX4* p.Tyr186X hiPSCs and from isogenic BLCs homozygous for p.X186 or p.Tyr186X allele were smaller but directionally consistent (Fig. 7e and Supplementary Table 6), supporting p.X186 (causing *PAX4* haploinsufficiency) resulted in dysregulated gene expression. The glycolysis stress test revealed a rescue in glycolytic reserve only in cells corrected for the p.His192 allele (Extended Data Fig. 7a-b) but not for cells corrected for the p.Tyr186X allele (Extended Data Fig. 7c-d). Importantly, correcting the *PAX4* p.His192His and p.Tyr186X mutations significantly increased and restored the total insulin content of the BLCs (Fig. 7f-g), indicating that the *PAX4* variants were a direct cause of reduced insulin content in the donor-derived BLCs.

## Discussion

Rodent models have demonstrated that *Pax4* plays an important role in beta cell specification during early pancreas development^13,40^. However, differences between rodent and human islets in architecture^41^ and gene expression^42^ make it challenging to extrapolate data based on rodent studies directly to humans. For instance, heterozygous *Pax4* knockout mice do not develop diabetes^13^ but *PAX4* variants causing altered transcriptional activity are strongly associated with increased diabetes risk in humans^8,16,43,44^. These observations suggest that human beta cells could be more sensitive to changes in *PAX4* gene dosage.

While *PAX4* p.Arg192His has been identified as one of the most reproducible variants uniquely associated with East Asian T2D, the role of *PAX4* or its variant p.Arg192His in human beta cell development has not been addressed. Our study capitalized on access to East Asian carriers of T2D *PAX4* risk alleles to study their effect on human beta cell function *in vivo*, and generate donor-derived hiPSCs as a versatile platform to interrogate the role of *PAX4* during human pancreas development *in vitro*^45–48^. Our clinical phenotyping of the *PAX4* p.Arg192His allele carriers demonstrated decreased pancreatic beta cell function based on AIRg, HOMA-B, and lowered DI despite the donors being insulin sensitive based on HOMA-IR measures. Whilst our investigation of the impact of a novel variant predicted to result in a loss of PAX4 function (p.Tyr186X) demonstrated that *PAX4* haploinsufficiency is insufficient to cause monogenic diabetes, it was consistent with a negative impact on beta cell function. This finding is further supported by large sequencing studies that collectively show an association of rare alleles in *PAX4* with an increased risk for diabetes or elevated HbA1c levels.

While donor hiPSC-derived beta cells can be used to study human pancreas development *in vitro*, this experimental model suffers from the following challenges: 1) hiPSC line-to-line variability, 2) the heterogenous nature of differentiated islet-like cells and 3) incomplete functional maturity of differentiated beta-like cells^49^. To circumvent these challenges while leveraging the benefits of this model, we rigorously applied two differentiation protocols to multiple donor-derived and isogenic genome-edited hiPSC lines. Our three independent sets of RNA-seq data using two protocols^22,25^ in multiple hiPSC models (*PAX4*^-/-^ knockout, donor-derived and isogenic hiPSCs carrying *PAX4* variants, and donor-derived gene-corrected hiPSCs)^27^ demonstrated that all differentiating cells shared a similar trajectory towards pancreatic islet-like cells. Knockout of *PAX4* did not result in the ablation of human beta cells, but rather, resulted in compromised beta cells with elevated expression of multiple endocrine hormone markers and lowered expression of genes associated with beta cell functional maturation. These observations were similarly replicated across donor-derived hiPSCs of three independent genotypes (p.Arg192His, p.His192His and p.Tyr186X), whereby BLCs carrying *PAX4* alleles demonstrated increased polyhormonal gene expression and reduced total insulin content. Our molecular assessments confirmed p.X186 to undergo NMD, while the p.His192 resulted in altered transcriptional activity. Contrary to rodent models, our human *in vivo* and *in vitro* findings indicate that differentiating human beta cells are sensitive to the functional (haploinsufficiency; loss-of-function) *PAX4* gene dosage required to maintain beta cell identity, insulin production and secretion. Our data are consistent with a recent study on *HNF1A* deficiency^50^ and support a model where *PAX4* T2D-risk alleles mediate disease risk by biasing endocrine precursor cells towards an alpha cell fate.

Transcriptomic assessment of EPs identified altered metabolic signatures in carriers of p.Arg192His or p.Tyr186X allele(s). Indeed, metabolic stress can compromise beta cell identity and has been proposed to be one of the mechanisms underlying beta cell exhaustion in T2D^51,52^. Unfortunately, the use of NAC to alleviate oxidative stress was insufficient to rescue the total insulin content in *PAX4* variant-expressing BLCs. BLCs derived from gene-corrected hiPSCs demonstrated a rescue in total insulin content, affirming the direct contribution of p.Arg192His or p.Tyr186X to decreased insulin content. While we were unable to determine the direct cause of the altered metabolic signature observed in EPs or whether the variants were a direct cause of the metabolic signature, it is tempting to hypothesize that the inferior beta cells resulting from the *PAX4* variants, compounded with cellular metabolic stress, hasten the eventual progression toward T2D development.

As our hiPSC-derived beta cells were not functional *in vitro*, we included the study of *PAX4* in the human beta cell line EndoC-βH1. Knockdown of *PAX4* led to a derepression of the *GCG* gene promoter and elevated *GCG* gene expression in beta cells. *PAX4* expression requires cooperative activation by several key transcription factors specific to pancreatic beta cells^12^ to specify PAX4 exclusively in beta cells. In rodents, the maintenance of pancreatic beta cell identity requires a continual repression of non-beta cell gene expression^51,52^. The expression of multiple hormonal markers, including GCG, is one of the hallmarks of immature cells that could have diminished function in endocrine hormone secretion. The reduced *PAX4* levels resulting in the loss-of-repression of non-beta cell gene expression could possibly explain the co-expression of GCG in C-PEP-expressing BLCs carrying the p.His192 or p.X186 allele(s). We have observed that carriers of p.Arg192His or p.Tyr186X alleles secrete less insulin during GSIS (Fig. 1a and f), and this was recapitulated in our si*PAX4* and sh*PAX4* EndoC-βH1 cells (Fig. 2). The insulin content within pancreatic islet cells has a strong correlation with the amount of insulin secreted during GSIS^53^. The reduced total insulin content observed in our hiPSC and EndoC-βH1 models and subsequent impaired GSIS in EndoC-βH1 cells collectively suggest a role for PAX4 in maintaining beta cell identity and regulating insulin secretion function.

A limitation of our study is the description of a single family with a *PAX4* PTV, limiting the confidence with which conclusions can be drawn from our observations of an effect of *PAX4* haploinsufficiency in humans. We sought to strengthen our findings through aggregation of exome-sequencing data from multiple publicly available datasets, which both provided nominal evidence for a role of rare coding variation in elevated diabetes risk and support mounting evidence that *PAX4* is not a monogenic diabetes gene^10^.

The loss of beta cell identity and the acquisition of polyhormonal cells have been reported in the pancreatic islets of individuals with diabetes^54,55^. The transdifferentiation of metabolically stressed beta cells to express GCG has been proposed by several groups as a mechanism underlying beta cell failure in T2D^55–57^. In the current study, we demonstrate how coding gene variants in *PAX4* can influence pancreatic beta cell development, identity, and function, thereby predisposing East Asian carriers to higher risks of developing T2D.

## Methods

### Clinical studies

We recruited 183 non-diabetic individuals by genotype (62 carriers of p.Arg192His allele and 121 controls) from existing research programs in Singapore. The inclusion criteria were Chinese ethnicity, age between 21 and 80 years, non-smoker or no use of nicotine or nicotine-containing products for at least 6 months. Subjects with a known history of diabetes mellitus, screening HbA1c greater than 6.5% or fasting plasma glucose greater than 7.0 mmol/L were excluded. Subjects with weight loss greater than 5% of body weight in the preceding six months, major surgery in the last three months, a history of malignancy, estimated creatinine clearance based on the MDRD formula less than 60 mL/min, current corticosteroid use, or any clinically significant endocrine, gastrointestinal, cardiovascular, hematological, hepatic, renal, respiratory disease, or pregnancy were also excluded. The study was approved by the National Healthcare Group Domain Specific Review Board (2013/00937), and informed consent was obtained from all participants. Data on demographics and medical history were obtained through an interviewer-administered questionnaire. Height and weight were measured. All subjects underwent confirmation of genotype by polymerase chain reaction-restriction fragment-length polymorphism analysis.

### Frequently sampled intravenous glucose tolerance test

A 3-hour intravenous glucose tolerance test was performed after an overnight 10- to 12- hour fast. Subjects were required to abstain from strenuous physical activity, alcohol and caffeinated beverages 24 hours before the procedure. A bolus of intravenous 50% glucose (0.3 g/kg body weight) was given within 60 seconds into the antecubital vein. Regular insulin (Actrapid; NovoNordisk, Copenhagen, Denmark) was administered as a bolus injection at 20 min at a dose of 0.03 units/kg body weight. Blood was sampled from the contralateral antecubital vein at -15, -10, -5, 0, 2, 3, 4, 5, 6, 8, 10, 14, 19, 22, 25, 30, 40, 50, 70, 100, 140 and 180 min for assessment of plasma glucose (YSI 2300 STATPLUS; YSI Incorporated, Life Sciences, Yellow Springs, OH, USA) and insulin (Advia Centaur; Siemens Health-care Diagnostics, Hamburg, Germany). AIRg (acute insulin response to glucose) and Si (insulin sensitivity) were estimated using mathematical modeling methods (MINMOD Millennium, ver. 6.02) Disposition index (DI) was calculated as AIRg x Si.

### Oral glucose tolerance test (OGTT)

Fifty-seven subjects (29 heterozygous p.Arg192His carriers and 28 p.Arg192Arg controls) were invited to return for a 3-hour oral glucose tolerance test. The test was performed after an overnight 10 to 12-hour fast. A 75-gram glucose drink in 200 mLs of water was administered orally over 5 min. Blood samples were collected via an intravenous cannula at -10, 0, 10, 20, 30, 45, 60, 75, 90,120, 150 and 180 min for glucose (YSI 2300 STATPLUS; YSI Incorporated, Life Sciences, Yellow Springs, OH, USA), insulin (Advia Centaur; Siemens Health-care Diagnostics, Hamburg, Germany), glucagon (Human Glucagon ELISA; BioVendor R&D, Shizuoka, Japan) and GLP-1 (Glucagon-like peptide-1 total ELISA; IBL International, Hamburg, Germany). HOMA-B was calculated using the formula: 20 × fasting insulin (μIU/mL)/fasting glucose (mmol/mL) − 3.5. HOMA-IR was computed using the formula: fasting insulin (μIU/ml) × fasting glucose (mmol/mL)/ 22.5.

### Statistical analysis of clinical data

Analyses were carried out using SPSS software version 18.0 (SPSS Inc., Chicago, IL, USA). Independent t-test and Chi-square tests were used to compare continuous and categorical variables between carriers and controls, respectively. A multiple linear regression model was used with adjustment by age, sex and BMI. Data are shown as the means (SD), and a p-value of <0.05 was considered statistically significant.

### Cell culture

The use of human cells is covered by A*STAR IRB 2020-096. All mammalian cells were routinely tested to be mycoplasma free using a MycoAlert^TM^ PLUS mycoplasma detection kit (Lonza Bioscience, LT07-710). All mammalian cells were cultured in a 5% CO_2_ humidified incubator at 37°C. Unless otherwise stated, cells were passaged using 0.25% trypsin. Mouse insulinoma 6 (MIN6) cells were cultured in high glucose DMEM (HyClone, SH30021.01) supplemented with 15% FBS (HyClone, SV30160.03), 1% sodium pyruvate (ThermoFisher Scientific, 11360070) and 55 µM beta mercaptoethanol (Gibco, 21985-023). Alpha TC clone 9 mouse pancreatic adenoma cells (αTC1.9) (ATCC, CRL-2350™) were cultured in low glucose DMEM (1.0 g/L) supplemented with an additional 1.0 g/L glucose (final glucose concentration to be 2.0 g/L), 10% FBS, 15 mM HEPES (ThermoFisher Scientific, 15630080), 1% NEAA (Gibco, 11140-50), 0.02% Bovine serum albumin (BSA) (Sigma-Aldrich, A9418) and 1.5 g/L sodium bicarbonate (ThermoFisher Scientific, 25080094). AD293 (Agilent, 240085) and 293FT (Invitrogen, R70007) human embryonic kidney cell lines were cultured in high glucose DMEM supplemented with 10% FBS and 1% NEAA. EndoC-βH1 cells^58^ (Human Cell Design) were cultured according to the manufacturer’s recommendations. Briefly, tissue culture plates were precoated with high glucose DMEM supplemented with 2 µg/mL fibronectin (Sigma-Aldrich, F1141) and 1% ECM (Sigma-Aldrich, E1270) at least 30 min prior to cell plating. Low glucose DMEM (Gibco, 11885084) supplemented with 2% BSA, 10 mM nicotinamide (Sigma-Aldrich, N0636 or N3376), 2 mM GlutaMAX^TM^ (Gibco, 35050061), 50 µM beta mercaptoethanol, 5.5 μg/mL transferrin (Sigma-Aldrich, T8158) and 6.6 ng/mL sodium selenite (Sigma-Aldrich, 214485). Cells were passaged weekly with 0.05% or 0.25% Trypsin and neutralized with 20% FBS in DPBS and plated at a density of 70,000 cells/cm^2^. The isogenic SB Ad3.1 hiPSC line derived from human skin fibroblasts from a Caucasian donor with no reported diabetes (Lonza CC-2511, tissue acquisition number 23447) was obtained from the Human Biomaterials Resource Centre, University of Birmingham. The SB line and donor-derived hiPSC lines generated herein were cultured in TeSR^TM^-E8^TM^ or mTeSR-1^TM^ medium (StemCell Technologies, 05990 or 85850) with daily media changes. hiPSCs were passaged twice weekly using ReLeSR^TM^ (StemCell Technologies, 05872) or Accutase (Gibco, A1110501) according to the manufacturer’s instructions. Culture plates were precoated with 0.1% gelatin in cell culture grade water for at least 10 min and then with MEF media for at least 48 hours prior to plating or with Corning Matrigel hESC-Qualified Matrix (VWR International, BD354277) for at least an hour prior to plating.

### Generating donor-derived hiPSC lines

Skin punch biopsies were obtained from the upper forearm of recruited subjects and cultured in low glucose DMEM supplemented with 10% heat-inactivated FBS and 1% MEM non-essential amino acids (Gibco, 11140-50) to obtain fibroblasts. A Human Dermal Fibroblast Nucleofector™ Kit (Lonza Bioscience, VDP-1001) was used for episomal reprogramming of fibroblasts. Cells were trypsinized and washed with DPBS and 500,000 cells were resuspended in Nucleofector^TM^ Solution according to manufacturer’s instructions. The following Yamanaka factors from Addgene were added at 1 µg to the cell suspension: pCXLE-hOCT3/4-shp53-F (plasmid #27077), pCXLE-hSK (plasmid #27078), and pCXLE-hUL (plasmid #27080). Nucleofection program P22 was used to transfect cells. At the end of the nucleofection, cells were plated onto mitotically-inactivated CF1-MEF (plated one day in advance) (Lonza Bioscience, GSC-6201G) and cultured in DMEM/F12 (Gibco, 10565018) media supplemented with 20% KnockOut^TM^ serum replacement (Gibco, 10828010), 1% NEAA, and 10 ng/mL FGF-2 (Miltenyi Biotec, 130-093-842). Media were replaced daily until hiPSC colonies emerged.

Peripheral blood mononuclear cells (PBMCs) were extracted from donor blood using a BD Vacutainer® CPT™ Mononuclear Cell Preparation Tube (BD Biosciences, 362753). The white buffy coat layer containing PBMCs was collected, washed twice with DPBS, and centrifuged at 300xg for 10 min to pellet the cells. One to two million cells were seeded and cultured in expansion media: IMDM media (Gibco, 12440053) supplemented with 10% FBS, 50 µg/mL of L-ascorbic acid (Sigma-Aldrich, A8960), 50 ng/mL of Stem Cell Factor (RnD Systems, 255-SC-010), 10 ng/mL IL-3 (StemCell Technologies, 78040), 2 U/mL Erythropoietin, 40 ng/mL IGF-1 (BioVision, 4119), 1 µM dexamethasone (Sigma-Aldrich, D8893) and 0.2% Primocin (Invivogen, ant-pm-1). PBMCs were reprogrammed following manufacturer’s instructions using CytoTune™-iPS 2.0 Sendai Reprogramming Kit (Invitrogen™, A16517) to obtain hiPSCs. For reprogramming, 200,000 PBMCs were plated in 12-well plates, and Sendai viruses [hKOS (MOI5), hc-Myc (MOI5), and hKlf4 (MOI3)] were added to culture media supplemented with 8 μg/mL of Polybrene (Sigma-Aldrich, TR-1003). Media was replaced the next day. Cells were collected and plated onto mitotically-inactivated CF1-MEFs (pre-seeded one day in advance) and cultured in DMEM/F12 media supplemented with 20% KOSR (Gibco, 10828010), 1% NEAA (Gibco, 11140-50) and 10 ng/ml FGF-2 (Miltenyi Biotec, 130-093-842), supplemented with 50 µg/mL of L-ascorbic acid, 50 ng/mL of Stem Cell Factor, 10 ng/mL IL-3, 2 U/mL Erythropoietin, 40 ng/mL IGF-1, and 1 µM dexamethasone for the first two days. The reprogrammed PBMCs were then maintained in basal media without growth factor and small molecule supplementation until hiPSC colony formation. hiPSC colonies were handpicked and cultured with a TeSR^TM^-E8^TM^ Kit. Each colony was designated to be one hiPSC line and expanded for cryopreservation, with two to three independent lines per donor.

Immunofluorescence staining was performed on all hiPSC lines used in this study to confirm the expression of pluripotency markers OCT3/4, SOX2, NANOG, SSEA-4 and TRA1-60. One representative donor-derived hiPSC line from each donor was submitted for karyotyping (Cytogenetic laboratories, Singapore General Hospital) and for teratoma assay (A*STAR Biological Resource Centre (BRC) Animal Facility). Teratomas were then sent to Advanced Molecular Pathology Laboratory (AMPL, A*STAR) for paraffin block processing, sectioning and H&E staining. Derivation of all three germ layers (definitive endoderm, mesoderm and ectoderm) was confirmed using light microscopy.

### CRISPR-Cas9 genome editing of hiPSCs

To generate *PAX4*^-/-^ isogenic SB Ad3.1 hiPSC lines, a strategy was designed to mirror the well-studied *Pax4*^-/-^ mice where almost all of the functional domains were replaced with a beta galactosidase-neomycin resistance cassette^13^. To delete the majority of the paired and homeodomains, sgRNAs were designed targeting exon 2 (sgRNA#1: CTAGGGCGTTACTACCGCAC) and exon 5 (sgRNA#2: TATCCTGATTCAGTGGCCCG) of *PAX4* gene (ENST00000341640.6). To generate the p.Arg192His and p.Tyr186X variants in the SB Ad3.1 hiPSC line, sgRNA#2 was electroporated with HDR template with either rs2233580 (G>A) mutation or GTA duplication, respectively. To correct the donor-derived p.His192His and p.Tyr186X hiPSCs, sgRNA#3 (GGCAGTAGCCAGCTTTCCAT) or sgRNA#4 (ATCTCCGCAGAGTTCCAGCG) were electroporated with an HDR repair template. sgRNAs were synthesized following manufacturer’s instructions using the EnGen sgRNA Synthesis Kit, *S. Pyogenes* (NEB, E3322), followed by DNase treatment and RNA purification using the RNA Clean & Concentrator Kit (Zymo Research, R1017). Ribonucleoprotein (RNP) complexes were formed by combining 20 μM (681 ng) sgRNA, 20 μM Cas9 (NEB, M0646T) and Buffer R (ThermoFisher, MPK109R) in a total volume of 6 μL and incubating at room temperature for 15 min. The RNP complex was then combined with 250,000 hiPSCs in 15 μL of Buffer R and incubated on ice for 5 min. Ten microliters of the RNP+cell mixture was electroporated in two separate electroporations using the Neon™ Transfection System 10 μL Kit (ThermoFisher, MPK1025). Electroporated cells were seeded into Matrigel-coated plates with mTeSR media and 10 μM Y-27632 (StemCell Technologies, 72302). Forty-eight hours after electroporation, hiPSCs were plated at low-density (5,000 cells/60 mm dish) on Matrigel-coated plates with mTeSR and 10 μM Y-27632. The resulting colonies were handpicked and expanded for further genotyping and quality control measures.

### hiPSC differentiation to into BLCs

For differentiation experiments using Protocol A, hiPSCs were cultured in mTeSR™1 with daily media changes and passaged using Accutase. Cells were plated at 10^6^ cells/well in Growth Factor Reduced Matrigel (Corning, 356230)-coated CellBind 12-well tissue culture plates (Corning, 356230 and 3336) in mTeSR1 (StemCell Technologies, 85850) supplemented with 10 μM of Y-27632 dihydrochloride (AbCam, ab120129). The following morning, the medium was changed to mTeSR™1, and differentiation was started 24 hours after plating. Directed differentiation protocol was adapted from Rezania et al. and basal differentiation media (using MCDB-131) was formulated accordingly^22^. Media was changed daily to basal media supplemented with growth factors and small molecules (Table 1) with the following modifications: Activin A and CHIR 99021 were used for Stage 1; all stages were performed in planar culture; and stages 6 and 7 were both 6 days in length.

**Reagent Table 1:** directed differentiation protocols A and B.

| **Reagent** | **Source, Catalog number** |
| --- | --- |
| MCDB 131 media | Corning, 15-100-CV; Gibco, 10372-019 |
| CMRL-1064 supplemented | Mediatech, 99-663-CV |
| Bovine serum albumin | Roche, 10775835001; Proliant, 68700 |
| Insulin-Transferrin-Selenium-Ethanolamine | Gibco, 51500-056 |
| L-Ascorbic acid | Sigma-Aldrich, A4544 or A8960 |
| Heparin | Sigma-Aldrich, H3149 |
| Human recombinant Activin A | Peprtech, 120-14; StemCell Technologies, 78001.2 |
| CHIR 99021 | Axon Medchem, 1386; Tocris, 4423 |
| KGF/FGF-7 | Peprtech, 100-19; StemCell Technologies, 78046.2 |
| Retinoic acid | Sigma-Aldrich, R2625; WAKO, 186-01114 |
| SANT1 | Sigma-Aldrich, S4572; Santa Cruz, sc-203253 |
| Phorbol 12,13-dibutyrate | Tocris, 4153 |
| LDN-193189 | Stemgent, 04-0074; Sigma-Aldrich, SML0559 |
| Compound E, γ-secretase inhibitor XXI | EMD Millipore, 565789; Cayman, 15579-20 |
| Alk5 inhibitor II | ENZO, ALX-270-445-M001 |
| L-3,3',5-triiodothyronine | Sigma-Aldrich, T6397; Merck Millipore, 642511 |
| Human betacellulin | Cell Signalling, 5235SF |
| Alpha-Amyloid Precursor Protein Modulator | EMD Millipore, 565740 |
| N-acetyl cysteine | Sigma-Aldrich, A9165 |
| R428 | SelleckChem, S2841 |
| Trolox | EMD Millipore, 648471 |
| Zinc Sulfate | Sigma-Aldrich, Z0251 |

For differentiation experiments using Protocol B, hiPSCs were plated and maintained in 10 cm plates until 80-90% confluency. Following, hiPSCs were washed with DPBS and dissociated into single cells using TrypLE™ Express (Gibco, 12605-010). Cells were seeded at a density of 1 million cells per mL of mTeSR™1 kit supplemented with 10 μM of Y-27632 (StemCell Technologies, 72303) on non-treated 6 well plate. Cells were then incubated in tissue culture incubator on an orbital shaker with a shaking speed set at 80 rpm over a duration of 24 – 48 hours before the start of differentiation by changing to differentiation media supplemented with growth factors and/or small molecules. Directed differentiation protocol was adapted from Pagliuca et al. and basal differentiation media (S1, S2, S3, S5 and S6) were formulated accordingly^25^. Fresh differentiation media supplemented with growth factors and small molecules were added at stipulated timepoints for directed differentiation over a duration of 35 days. The details of the culture medium and key reagents used for Protocol B can be found in Table 1.

### Cloning

*PAX4* plasmid (pLenti6.2/V5-DEST-PAX4, HsCD00329734) was purchased from DNASU plasmid repository. Full-length *PAX4* sequence was confirmed via Sanger sequencing before subcloning into an engineered lentiviral vector pCDH-MCS-EF1-GFP to include a 5’ Flag tag and a 3’ V5 tag within the multiple cloning site (MCS). The full-length *PAX4* sequence was subcloned into the pCDH-MCS-EF1-GFP vector for protein expression. Refer to Table 2 for the list of primers used for cloning in this study. Cloning primers h*Pax4*FLXbaI1F (forward) and h*Pax4*FLV5Xho1R (reverse) were used to amplify full-length *PAX4* sequence. Polymerase chain reaction (PCR) was performed using Phusion™ High-Fidelity DNA Polymerase (ThermoScientific, F530) for sequence amplification. Thermal cycling conditions were set according to the manufacturer’s manual. Restriction enzyme digestion was performed on the pCDH-MCS-EF1-GFP and amplified *PAX4* sequence independently using XbaI (New England Biolabs, R0145) and XhoI (New England Biolabs, R0146) according to the manufacturer’s manual. Digested products were resolved using gel electrophoresis and gel extraction was performed using Purelink^TM^ Quick Gel Extraction Kit (Invitrogen, K210012). Ligation was performed using Quick Ligation Kit (New England Biolabs, M2200S) according to the manufacturer’s manual. The ligated plasmids were transformed into home-made competent cells propagated from Stbl3™ competent cells (Invitrogen, C7373-03) and sequentially amplified in LB broth for plasmid extraction using PureLink™ HiPure Plasmid Filter Maxiprep Kit (Invitrogen, K210017). To introduce R192H and Y186X mutations into the pCDH-*PAX4* plasmid, site-directed mutagenesis (SDM) primers were designed (refer to Table 2) and SDM was performed according to the procedures described.

**Reagent Table 2:**
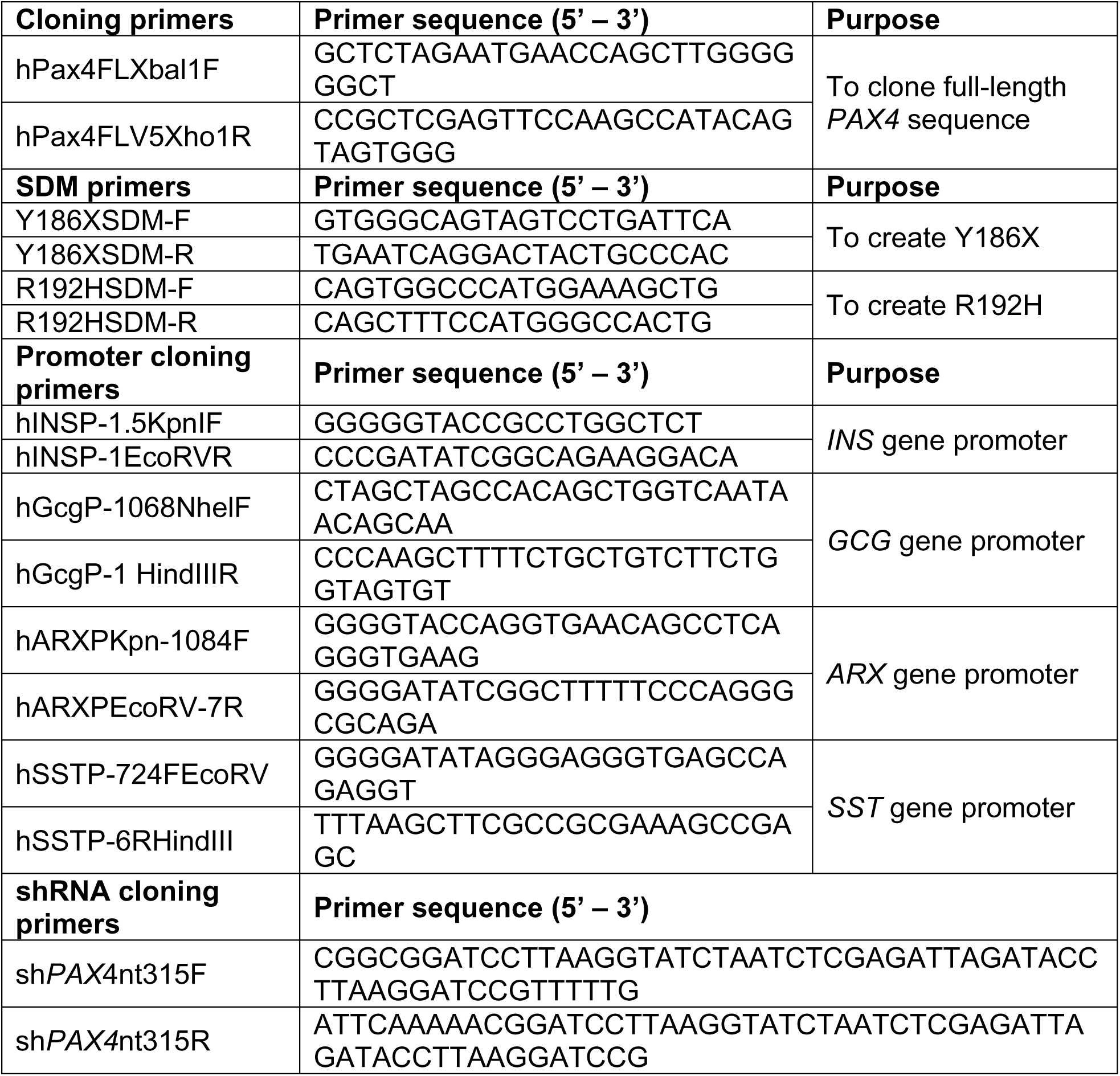
primers for cloning.

For gene promoter cloning, human genomic DNA extracted from AD293 cells was used as template. Basic luciferase vector, pGL4.10 (Promega) was used as cloning vector for gene promoters. Briefly, primers targeting the promoter region (−1 nucleotide from ATG translational start site) were designed (refer to Table 2). Targeted promoter regions were amplified, digested with restriction enzyme, ligated and transformed into competent cells similarly as described in the previous section. With the exception of the insulin gene promoter, all other gene promoters used in this study were amplified from human gDNA and subcloned into pGL4.10 at the multiple cloning site. The pGL4.10 *INS* promoter plasmid was synthesized by IDT (gBlocks™ Gene Fragments). Length of the various gene promoters used is as follows: *PAX4* – 1384 bp, *INS* – 1499 bp, *GCG* – 1068 bp, *SST* – 718 bp.

The design of shRNA sequence to knockdown *PAX4* gene was referenced to Genetic Perturbation Platform (Broad Institute), Clone ID TRCN0000015989. The shRNA targets the coding sequence CGGATCCTTAAGGTATCTAAT within *PAX4* gene (Table 2). The shRNA was ligated into pLKO.1 vector using Quick Ligation Kit (New England Biolabs, M2200S) according to the manufacturer’s instructions. Successfully ligated sh*PAX4* plasmid was amplified for subsequent experiments.

### Lentiviral-mediated sh*PAX4* stable line generation

3^rd^ generation lentivirus system was used for this study. Lentivirus plasmids used for virus production: pRC/CMV-Rev (Rev), pHDM-HIVgpm (Gag/Pol) and pHDM-G (Vsv-g). Non-targeting shScramble and shRNA targeting human *PAX4* (sh*PAX4*) gene were subcloned into pLKO.1 vector for lentiviral packaging in 293FT cells. For the generation of stable lines, EndoC-βH1 cells were plated onto 10 cm plates. The cells were then transduced with pLKO.1 shScramble or sh*PAX4* lentiviruses in the presence of 8 µg/mL polybrene. After 72 hours, the transduced cells were cultured in EndoC media supplemented with 500 μg/ml of G418 antibiotic (Invivogen, ant-gn-1). In parallel, one plate of untreated EndoC-βH1 cells (plated at the same density) was cultured in the same antibiotic supplemented media as a control. Media were replenished routinely during this selection process. Thereafter, the surviving EndoC-βH1 cells were expanded to obtain stable lines.

### EndoC-βH1 gene silencing using siRNAs

Knockdown studies in EndoC-βH1 cells were performed using Lipofectamine RNAiMAX^®^ transfection protocol and 15 nM SMART pool ON-TARGETplus siRNAs (Horizon Discovery Biosciences, si*NT*: D-001810-10-05, si*PAX4*: L-012240-00-0005) diluted in Opti-MEM reduced serum-free medium (ThermoFisher Scientific, 31985062) and 0.4% RNAiMAX^®^ (ThermoFisher Scientific, 13778150). Silencing efficiency was determined by qPCR from samples collected during GSIS, five days post-transfection.

### Glucose-stimulated insulin secretion (GSIS) assay

siNT and si*PAX4* EndoC-βH1 cells were seeded in 48-well plates at a density of 180,000 cells six days prior to GSIS. Cells were gently washed three times with pre-warmed secretion assay buffer (114 mM sodium chloride, 4.7 mM potassium chloride, 1.2 mM calcium chloride, 1.2 mM potassium phosphate, 1.16 mM magnesium sulphate, 25 mM sodium bicarbonate, 0.2% fatty acid-free BSA (Proliant, 68700), 20 mM HEPES, adjusted to pH 7.3). Cells were then incubated in secretion assay buffer for 1 hour before being stimulated with 2.8 mM or 16.7 mM glucose for 40 min. Supernatant was collected at the end of 40 min for insulin secretion measurements and insulin content was collected using RIPA buffer. AlphaLISA human insulin research kit (Perkin Elmer, AL204C) was used to measure insulin secretion and content. Total protein measurements were determined using a Pierce BCA Protein Assay Kit (Life Technologies, PI23227). Stimulation index was calculated by normalizing to total protein and relative to 2.8 mM glucose.

sh*PAX4* and shScramble EndoC-βH1 cells were seeded in 12-well plate prior to GSIS assay. Before GSIS assay, cells were gently washed three times with warm Krebs Ringer bicarbonate (KRB) buffer (125 mM sodium chloride, 4.74 mM potassium chloride, 1 mM calcium chloride, 1.2 mM potassium phosphate, 1.2 mM magnesium sulfate, 5 mM sodium bicarbonate, 0.1% fatty acid-free BSA, 25 mM HEPES, adjusted to pH 7.5 (±0.2) with 1 M sodium hydroxide). After that, cells were subjected to normalization at 2.8 mM glucose for 1 hour before being stimulated at 2.8 mM and 16.7 mM glucose for 30 min each sequentially. At the end of each stimulation step, KRB buffer was collected for human insulin ELISA (Mercodia, 10-1113-10). Stimulation index was computed by insulin secreted at 16.7 mM divided by insulin secreted at 2.8 mM glucose. Total insulin was extracted from each sample after the whole process of GSIS was completed.

### Total insulin content extraction

At the end of the 35-day directed differentiation using Protocol B, BLCs from each hiPSC line were handpicked, and 400 μl of acid/ethanol solution was added. Cells were vigorously vortexed and subjected to repeated pipetting to break up cell clumps. BLCs were then incubated at 4°C overnight before total insulin extraction. For EndoC-βH1 cells, 500,000 cells were seeded onto each well of a 12-well plate. At the end of the experiment, cells were washed thrice with DPBS before adding 500 μl of acid/ethanol solution to each well and incubated at 4°C overnight prior to insulin extraction. After overnight incubation, the insulin extracts were centrifuged at 1000 rpm for 5 min. The top aqueous layer containing insulin was collected and subjected to human insulin ELISA assay (Mercodia, 10-1113-10) while the bottom layer (containing cell pellet) was boiled to dryness at 80°C on a heat block. The dried cell pellet was resuspended in water for total DNA quantification. All data involving total insulin content quantification were normalized to total DNA. For donor hiPSC-derived BLCs, the average total insulin content extracted from various cell lines from each donor in one experiment is represented as a single data point on the graph. For CRISPR-edited cells, each data point represents the average of total insulin extracted from one cell line in one experiment. For EndoC-βH1 cells, each data point represents the average of total insulin extracted in independently sampled triplicates in one experiment.

### Gene expression analysis

Total RNA was extracted using MN NucleoSpin RNA Kit (Macherey-Nagel). RNA was quantified and reverse transcribed to cDNA using High-capacity cDNA reverse transcription kit (Applied Biosystems, 4368813). QPCR was performed using iTaq™ Universal SYBR ® Green Supermix (Bio-rad, 172-5124). Thermal cycling was performed using CFX384 Touch Real-Time PCR System (Bio-rad). Relative quantification of each gene expression was normalized to *ACTIN*, calculated by the 2^-ddCt^ method. The qPCR primers used in this study are summarized in Table 3.

**Reagent Table 3:**
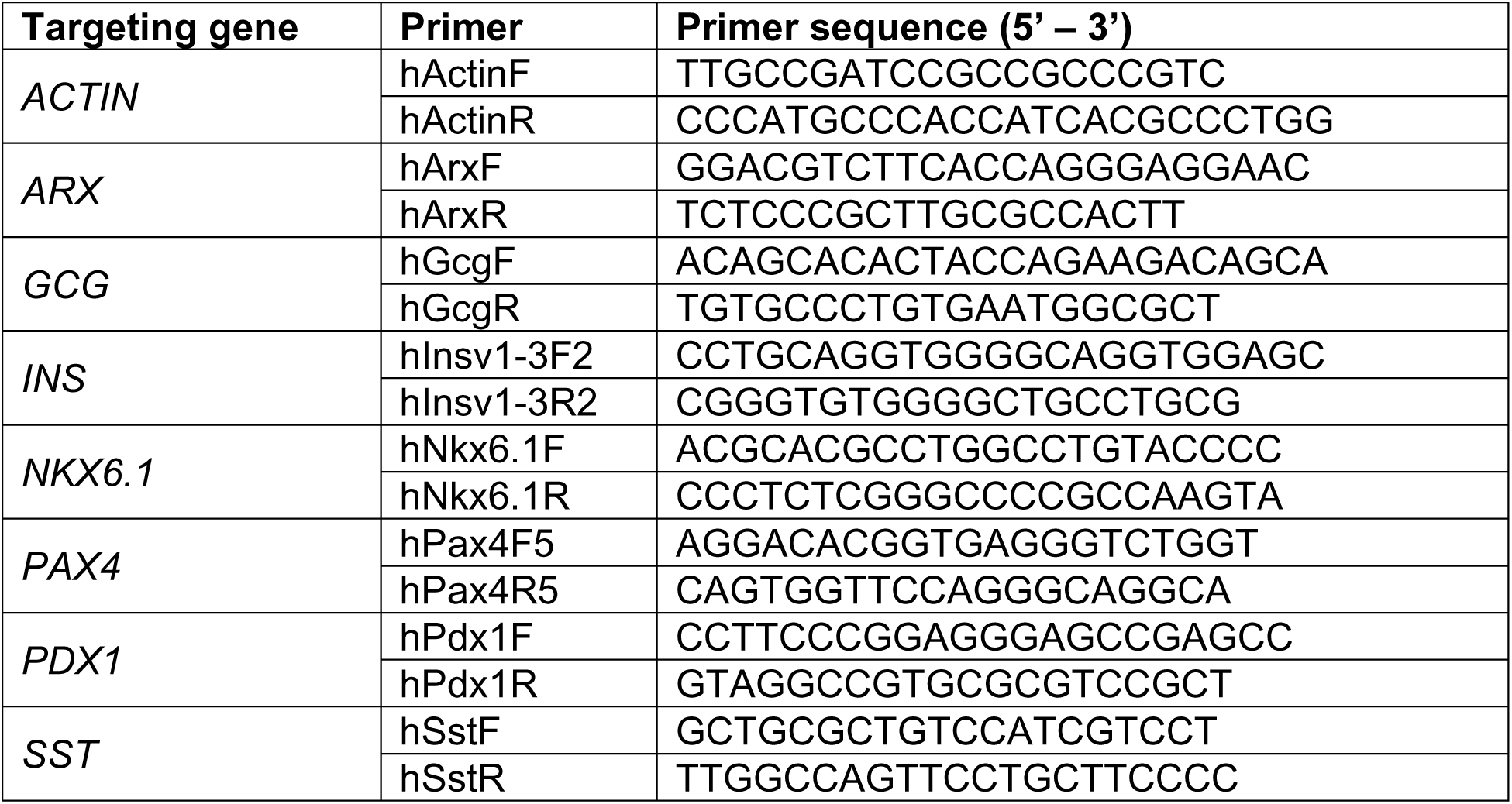
qPCR primers.

### Taqman allelic discrimination assay

Custom Taqman® Assay Design Tool (ThermoScientific) was used to design probes specific for either the wildtype (p.Arg192 or p.Tyr186) or *PAX4* variant transcripts (p.His192, ANT2HTM or p.X186, ANU7DDJ). A template sequence of approximately 600 bp around the SNP of interest was used as a reference to design custom assay probe. For Taqman assays qPCR, cDNA from EPs was used as templates. A 5 μl assay with TaqMan® SNP Genotyping MasterMix (Applied Biosystems, 4351384) was prepared according to the manufacturer’s manual. Relative fluorescence units (RFUs) from the HEX probe (wildtype allele) and the FAM probe (either Arg192 or X186 allele) were analyzed using the CFX384 Touch Real-Time PCR System (Bio-rad).

### Immunofluorescence staining

In preparation for pluripotency IHC, each donor-derived hiPSC line was seeded onto a few wells of precoated 12-well plate. AD293 and EndoC-βH1 cells were seeded onto uncoated and coated coverslips in 12-well plates, respectively. For overexpression studies, transfection was performed on seeded cells using Lipofectamine^TM^ 2000 transfection reagent (Invitrogen, 11668-019) or FuGENE® 6 transfection reagent (Promega, E2691) according to the manufacturer’s instructions. hiPSC-derived EPs and BLCs were collected and sent to Advanced Molecular Pathology Laboratory (AMPL, A*STAR) for cryo-embedding, cryo-block processing and sectioning. For IHC, cryosections were thawed and dried at room temperature before staining. Cells were washed thrice with DPBS and fixed with 4% paraformaldehyde (WAKO, 163-20145) for 20 min. Blocking and cell membrane permeabilization were performed using DPBS supplemented with 5% Donkey serum (Merck Millipore, S-30) and 0.1% Triton-X-100 (Merck Millipore, 9410) for 1 hour at 4°C. Cells were incubated with primary antibodies overnight at 4°C. Cryosections were then incubated with corresponding secondary antibodies for 1 hour at room temperature. For nuclear staining, cryosections were incubated with DAPI (1:5000) (Sigma-Aldrich, D9542) in DPBS for 20 min before mounting onto glass slides for imaging with Olympus Fluoview Inverted Confocal microscope. Refer to Table 4 for the list of antibodies used and their respective dilution factors for IHC.

**Reagent Table 4:** antibodies.

| **Antibody** | **Dilution factor** | **Cat#** | **RRID** |
| --- | --- | --- | --- |
| **Primary antibody** |  |  |  |
| Anti-beta Actin (Mouse monoclonal) | WB 1:10000 | Sigma-Aldrich, A5441 | AB_476744 |
| Anti-C-PEPTIDE (Rat monoclonal) | IF 1:100 | DSHB, GN-ID4 | AB_2255626 |
| Anti-FLAG M2 (Mouse monoclonal) | WB 1:1000; IF 1:100 | Sigma-Aldrich, F1804 | AB_262044 |
| Anti-Glucagon Antibody (N-17) (Goat polyclonal) | IF 1:100 | Santa cruz, sc-7780 | AB_641025 |
| Anti-Insulin antibody (Guinea pig polyclonal) | IF 1:100; FC 1:100 | Abcam, ab7842 | AB_306130 |
| Anti-PAX4 (Goat polyclonal)<br>*Not validated to stain endogenous protein expression | IF 1:100; WB 1:1000 | RnD, AF2614 | AB_2159529 |
| Anti-SOX2 antibody (Rabbit polyclonal) | IF 1:200 | Abcam, ab97959 | AB_2341193 |
| Anti-Human SSEA-4 Antibody, Clone MC-813-70 (Mouse monoclonal) | IF 1:100 | StemCell Technologies, 60062AD | AB_528477 |
| Somatostatin Antibody (D-20) (Goat polyclonal) | IF 1:100 | Santa cruz, sc-7819 | AB_2302603 |
| Anti-Human TRA-1-60 Antibody, Clone TRA-1-60R (Mouse monoclonal) | IF 1:100 | StemCell Technologies, 60064AD | AB_2686905 |
| Anti-V5 tag antibody (SV5-Pk1) (Mouse monoclonal) | IF 1:100; WB 1:1000 | Abcam, ab27671 | AB_471093 |
| Anti-CXCR4 PE-conjugated (Mouse monoclonal IgG<sup>2B</sup> Clone #44717) | FC 1:10 | RnD, FAB173P | AB_357083 |
| FC block or anti-mouse CD16/CD32 (Rat monoclonal) | FC 1:500 | BD, 553141 | AB_394656 |
| Alexa Fluor® 488 anti-Human Sox17 (Mouse monoclonal Clone #P7-969) | FC 1:20 | BD, 562205 | AB_10893402 |
| **Fluorophore-conjugated secondary antibody** |  |  |  |
| Alexa Fluor® 488 anti-goat | IF 1:500; FC 1:2000 | Invitrogen, A11055 | AB_2534102 |
| Alexa Fluor® 488 anti-mouse | IF 1:500; FC 1:2000 | Invitrogen, A21202 | AB_141607 |
| Alexa Fluor® 488 anti-rabbit | IF 1:500; FC 1:2000 | Invitrogen, A21206 | AB_2535792 |
| Alexa Fluor® 488 anti-rat | IF 1:500; FC 1:2000 | Invitrogen, A21470 | AB_10561519 |
| Alexa Fluor® 594 anti-goat | IF 1:500; FC 1:2000 | Invitrogen, A11058 | AB_2534105 |
| Alexa Fluor® 594 anti-Guinea pig | IF 1:500; FC 1:2000 | Invitrogen, A11076 | AB_141930 |
| Alexa Fluor® 594 anti-mouse | IF 1:500; FC 1:2000 | Invitrogen, A21203 | AB_141633 |
| Alexa Fluor® 594 anti-rabbit | IF 1:500; FC 1:2000 | Invitrogen, A21207 | AB_141637 |
| Alexa Fluor® 647 anti-Guinea pig | IF 1:500; FC 1:2000 | Jackson, 706-605-148 | AB_2340476 |
| Alexa Fluor® 647 anti-mouse | IF 1:500; FC 1:2000 | Jackson, A31571 | AB_162542 |
| Alexa Fluor® 647 anti-rabbit | IF 1:500; FC 1:2000 | Invitrogen, A31573 | AB_2536183 |
| Alexa Fluor® 488 Isotype | IF 1:500; FC 1:2000 | BD Pharmingen, 565572 | AB_2869685 |
| **HRP-conjugated secondary antibody** |  |  |  |
| Donkey anti-goat IgG-HRP | WB 1:10000 | Santa cruz, sc-2020 | AB_631728 |
| Mouse anti-goat IgG-HRP | WB 1:5000 | Santa cruz, sc-2354 | AB_628490 |
| Goat anti rabbit IgG HRP | WB 1:10000 | Santa cruz, sc-2004 | AB_631746 |
| Goat anti-mouse IgG-HRP | WB 1:10000 | Santa cruz, sc-2005 | AB_631736 |
| Goat anti-mouse IgG-HRP | WB 1:10000 | Santa cruz, sc-2055 | AB_631738 |
| m-IgGk BP-HRP | WB 1:5000 | Santa cruz, sc-516102 | AB_2687626 |
| Mouse anti-rabbit IgG-HRP | WB 1:5000 | Santa cruz, sc-2357 | AB_628497 |

WB: Western blot; IF: Immunofluorescence; FC: Flow cytometry.

### SDS-PAGE and Western blot

Cells were washed with DPBS and lysed in M-PER^TM^ (Mammalian protein extraction reagent) (ThermoScientific, 78501) in the presence of protease inhibitor cocktail (Sigma-Aldrich, P8340), phosphatase-2 inhibitor (Sigma-Aldrich, P5726), and phosphatase-3 inhibitor (Sigma-Aldrich, P0044). Protein was quantified using Pierce^TM^ BCA protein assay kit (ThermoScientific, 23227) according to the manufacturer’s instructions before being separated with SDS-PAGE and transferred to PVDF membrane. Protein blots were first blocked with 5% milk in TBST (1X Tris-Buffered Saline, 0.1% Tween 20) for 1 hour before incubating with primary antibody for either 2 hours at room temperature or overnight at 4°C. Blots were washed and then incubated with the respective HRP-conjugated secondary antibody for 1 hour. Chemiluminescence signals were visualized after incubation with Super Signal^TM^ West Dura Extended Duration Substrate (ThermoScientific, 34076). Refer to Table 4 for the list of antibodies used and their respective dilution factors for western blotting.

### Flow cytometry

DE cells generated with Protocol A were collected at the end of Stage 1 using Accutase. For extracellular staining, cells were washed twice with 1X Flow Cytometry Staining Buffer (RnD, FC001). Cells were blocked in Flow Cytometry Staining Buffer with FC block for 5 min before adding human anti-CXCR4 antibody for 45 min. Cells were then washed twice with Flow Cytometry Staining Buffer. For intracellular staining, cells were fixed using BD CytoFix Buffer (BD Biosciences, 554655) for 20 min on ice before washing twice with PBS. Using BD Perm/Wash Buffer (BD Biosciences, 554723), fixed cells were permeabilized for 30 min on ice, washed three times, and human anti-SOX17 antibody was incubated for 1 hour at 4 °C before a final wash step in PBS. Stained cells were acquired on SH800 Cell Sorter (Sony) and data analysis was performed using FlowJo™ 10.6.0.

EPs and BLCs were collected on D20 and D35, respectively, following differentiation with Protocol B before being dissociated into single cells using TrypLE™ Express (Gibco, 12605-010). Cells were passed through a 40 µm cell strainer and single cells were fixed with 4% PFA for 20 min on ice. Antigen blocking and cell permeabilization were performed using DPBS supplemented with 5% FBS (HyClone, SV30160.03) and 0.1% Triton-X-100 for 30 min on ice. Cells were stained with primary antibodies for 1 hour at room temperature. The cells were then washed three times with DPBS and incubated with corresponding secondary antibodies for 1 hour at room temperature. Flow cytometry analyses were performed with BD® LSR II Flow Cytometer (BD Biosciences) and data was analyzed using FlowJo^TM^ software (BD Biosciences). Refer to Table 4 for the list of antibodies used and their respective dilution factors for flow cytometry.

### Luciferase assays

Cells were plated in triplicate one day prior to co-transfection with 0.5 µg of pCDH-overexpression constructs encoding *PAX4* or its variants, 0.4 µg of pGL4.10 luciferase vector and 10 ng of TK Renilla vector. Transfection was performed using either Lipofectamine 2000 (Invitrogen, 11668-019) or FuGENE® 6 transfection reagent (Promega, E2691). Cells were lysed with lysis buffer at the end of transfection (24 hours for AD293, 48 hours for MIN6/αTC1.9 and 72 hours for EndoC-βH1 cells). Luciferase assay was performed using Dual-Glo® Luciferase Reporter Assay Kit (Promega, E2920) following the manufacturer’s instructions. The luciferase firefly activity was normalized against the Renilla readings within each well to account for variation in transfection efficiency across replicate wells. Each triplicate was normalized to the mean of the pCDH-MCS-EF1-GFP-empty control.

### Seahorse metabolic assays

EPs on D19 of directed differentiation using Protocol B were dissociated into single cells using TrypLE™ Express (Gibco, 12605-010) before passing through 40 μm cell strainer. 80,000 or 120,000 cells were plated with S5 differentiation medium^25^ supplemented with 10 μM of Y-27632 onto pre-coated Seahorse microplate one day prior to analysis. The same number of cells were seeded across all cell lines within each experiment. On the day of glycolysis stress test (Agilent Seahorse XF Glycolysis Stress Test Kit), cells were washed with unbuffered serum-free assay medium (DMEM 5030, Sigma-Aldrich; supplemented with 2 mM L-glutamine). Following, the cells were incubated in assay medium in a non-CO_2_ incubator at 37°C for 1 – 2 hours before measurements were taken. Extracellular acidification rates (ECAR) were measured using Seahorse XFe96 analyzer (Seahorse Bioscience) at pre-set timings prior to and following sequential injections of 10 mM glucose, 1.5 μM oligomycin and 50 mM 2-deoxy-glucose (2-DG). The same number of cells was seeded across the various genotypes for individual experiments. Four to eight technical replicates were seeded for each cell line. Each data point on graph represents the average of all replicates from one cell line. For the Mito Stress Test (Agilent Seahorse XF Cell Mito Stress Kit), cells were washed with unbuffered serum-free assay medium supplemented with 20 mM glucose (keeping the glucose level consistent with S5 differentiation medium), 2 mM pyruvate and 2 mM L-glutamine prior to incubation in assay medium in a non-CO_2_ incubator at 37°C for 1 – 2 hours before analysis. Oxygen consumption rates (OCR) were measured using Seahorse XFe96 analyzer prior to and following sequential injections of 1.5 μM oligomycin, 1 μM FCCP and 0.5 μM rotenone/antimycin-A. Similarly, the same number of cells was seeded across the various genotypes for individual experiments. Four to eight technical replicates were seeded for each cell line. One data point on the graph represents the average of all replicates from one cell line.

### RNA sequencing and analysis

Total RNA was extracted from samples generated using differentiation Protocol A at the end of Stage 1 (DE), 4 (PE), 5 (EP), and 7 (BLC) using RNeasy Mini Kit (Qiagen, 74104) following manufacturer’s instructions. Polyadenylated transcripts were isolated using NEBNext PolyA mRNA Magnetic Isolation Module (New England Biolabs, E7490). Sequencing libraries were prepared using the NEBNext Ultra Directional RNA Library Kit with 12 cycles of PCR and custom 8 bp indexes (New England Biolabs, E7420). Libraries were multiplexed and sequenced on the Illumina NovaSeq 6000 as 150-nucleotide paired-end reads. Reads were mapped to human genome build hg19 (GRCh37) using STAR v.2.5^59^, with GENCODE v19 (https://www.gencodegenes.org/human/release_19.html) as the transcriptomic reference. featureCounts from the Subread package v1.5 (http://subread.sourceforge.net/) was used to perform gene-level quantification. Differential expression analysis was performed per stage using DESeq2^60^ comparing *PAX4*^+/+^ and *PAX4*^-/-^ cell lines. First, the model was fit using a likelihood ratio test with genotype as a factor of interest and experiment as a covariate. From this, genes not within the top 5000 most significant genes were used as an empirical control (affected only by unwanted experimental variation) and the estimated factor of unwanted variation (k=1) was calculated using the RUVg function from RUVSeq^61^. To identify differentially expressed genes, DESeq2 was performed using a likelihood ratio test and including the factor from RUV as a covariate along with the technical replicate (experiment). Significance was determined by padj <0.05. Sashimi, TPM, and volcano plots were generated using ggplot2.

Total RNA was extracted from samples generated using differentiation Protocol B on day 0 (hiPSC), day 13 (PP2), day 20 (EP) and day 35 (BLC). Poly-A mRNA (10 – 100 ng) was used to construct multiplexed strand-specific RNA-seq libraries (NEXTflexTM Rapid Directional RNA-SEQ Kit, dUTP-Based, v2). The quality of individual libraries was assessed and quantified using Agilent 2100 Bioanalyzer and Qubit 2.0 fluorometer before pooling for sequencing using a HiSeq 2000 (1×101 bp read). Prior to cluster formation, pooled libraries were quantified using the KAPA quantification kit (KAPA Biosystems). The processing of raw RNA sequencing data was performed in collaboration with Molecular Engineering Laboratory (A*STAR) to remove low quality sequence reads. Filtered read sequences were mapped onto human genome (hg19). Fragments per kilobase million (FPKM) was used to calculate differential expression between patient lines using DESeq2. Using UMAP (Uniform Manifold Approximation and Projection) and PCA (Principal Component Analysis) dimension reduction clustering for all four time-points, 11 out of 164 samples were classified as outliers and excluded from clustering analyses.

For the generation of PCA plot for EPs differentiated using protocol B, the TPM read counts for each gene within the transcriptome were first normalized to calculate a standardized score. To derive a standardized score, we applied the following formula: log_10_(((GOI’s TPM counts for sample of interest +1)/(average TPM counts for GOI across all samples) + 1), where GOI represents a gene of interest within the whole transcriptome. Using this standardized score, PCA analysis was performed in R via the prcomp() function. The ggplot2 package was used to plot the final PCA biplot based upon the PC1 and PC2 loadings obtained from prcomp(). Finally, the stat_ellipse() function was used to cluster the transcriptome of cells with the various *PAX4* genotypes based upon a confidence interval of 90%.

## Data availability

Protocol A *PAX4*^-/-^ RNA-seq data: EGAS00001006036

Protocol B RNA-seq data 1: Submission into GEO in progress under NCBI #22948635

Protocol B RNA-seq data 2: GSE202206 (secure token: epufowcsnjcvvgv)

## Statistical analysis

Statistical analyses were performed using GraphPad Prism version 9. Data are presented as the standard error of the mean (SEM). Unless otherwise specified, unpaired Student’s t tests were performed to compare the means of two groups, and one-way ANOVA was performed to compare the means among three or more groups. A p-value of less than 0.05 indicates statistical significance.

## Abbreviations

2-DG: 2-deoxyglucose
AIRg: acute insulin response to glucose
AUC: area under the curve
BLC: beta-like cells
CABG: coronary artery bypass grafting
CHX: cycloheximide
CRF: chronic renal failure
DE: definitive endoderm
DI: disposition index
DM: diabetes mellitus
DR: diabetic retinopathy
ECAR: extracellular acidification rate
EP: endocrine progenitor
GDM: gestational diabetes mellitus
GO: gene ontology
GSIS: glucose-stimulated insulin secretion
HbA1c: hemoglobin A1c
HDR: homology directed repair
hiPSC: human induced pluripotent stem cells
HOMA-IR: homeostatic model assessment of insulin resistance
IGT: impaired glucose tolerance
IHD: ischemic heart disease
NAC: N-acetylcysteine
NMD: nonsense mediated decay
OCR: oxygen consumption rate
OGTT: oral glucose tolerance test
PBMC: peripheral blood mononuclear cell
PCA: principal component analysis
PE: pancreatic endoderm
PGT: primitive gut tube
PF: posterior foregut
PPM: permanent pace-maker implantation
PP1: pancreatic progenitor 1
PP2: pancreatic progenitor 2
PTV: protein truncating variant
qPCR: quantitative real-time PCR
RFU: relative fluorescence units
sgRNAs: single guide RNAs
Si: insulin sensitivity
SKAT: Sequence Kernel Association Test
SNP: single nucleotide polymorphism
T2D: type 2 diabetes
TPM: transcripts per million
UMAP: Uniform Manifold Approximation and Projection
WT: wildtype

## Acknowledgments

The authors thank A/P Yee Joo Tan for her support with antibodies and Daniela Moralli (University of Oxford) for her support with hiPSC karyotyping. We thank the Oxford Genomics Centre at the Wellcome Centre for Human Genetics (funded by Wellcome Trust grant reference 203141/Z/16/Z for the generation and initial processing of the sequencing data. H.H.L. is supported by the Institute of Molecular and Cell Biology (IMCB) Scientific Staff Development Award (SSDA) for her part-time Ph.D. N.A.J. K. is supported by the Stanford Maternal and Child Health Research Institute Postdoctoral Fellowship. A.L.G. is a Wellcome Senior Fellow in Basic Biomedical Science. A.L.G. is funded by the Wellcome (200837) and National Institute of Diabetes and Digestive and Kidney Diseases (NIDDK) (U01-DK105535, U01-DK085545, UM1DK126185, U01DK123743, U24DK098085) and the Stanford Diabetes Research Center (NIDDK award P30DK116074). A.K.K.T. is supported by IMCB, A*STAR, Lee Foundation Grant SHTX/LFG/002/2018, FY2019 SingHealth Duke-NUS Surgery Academic Clinical Programme Research Support Programme Grant, Precision Medicine and Personalised Therapeutics Joint Research Grant 2019, the 2nd A*STAR-AMED Joint Grant Call 192B9002, HLTRP/2022/NUS-IMCB-02, Paris-NUS 2021-06-R/UP-NUS (ANR-18-IDEX-0001), OFIRG21jun-0097, CSASI21jun-0006 and MTCIRG21-0071.

## Author contributions

Conceptualization: E.S.T., A.L.G., A.K.K.T.

Data curation: M.P.A., H.S., A.J.

Formal Analysis: H.H.L., N.A.J.K., M.P.A., H.S., A.J., A.L.G., A.K.K.T.

Funding acquisition: E.S.T., A.L.G., A.K.K.T.

Investigation: H.H.L., N.A.J.K., F.A., J.W.C., J.A., S.G., B.C., S.H., A.N.S.T., D.G., S.L.K., A.L.G., A.K.K.T.

Methodology: H.H.L., N.A.J.K., F.A., M.P.A., J.A., A.L.G., A.K.K.T.

Project administration: E.S.T., A.L.G., A.K.K.T.

Resources: H.H.L., N.A.J.K., F.A., J.A., S.H., D.G., S.L.K., E.S.T., A.L.G., A.K.K.T.

Software: M.P.A., A.J.

Supervision: E.S.T., A.L.G., A.K.K.T.

Validation: H.H.L., N.A.J.K.

Visualization: H.H.L., N.A.J.K.

Writing – original draft: H.H.L., N.A.J.K., A.L.G., A.K.K.T.

Writing – review & editing: All authors approved

## Competing interest statement

A.L.G.’s spouse is an employee of Genentech and holds stock options in Roche. A.K.K.T. is a co-founder of BetaLife Pte Ltd.

## Extended Data Figures

**Extended Data Fig. 1.**
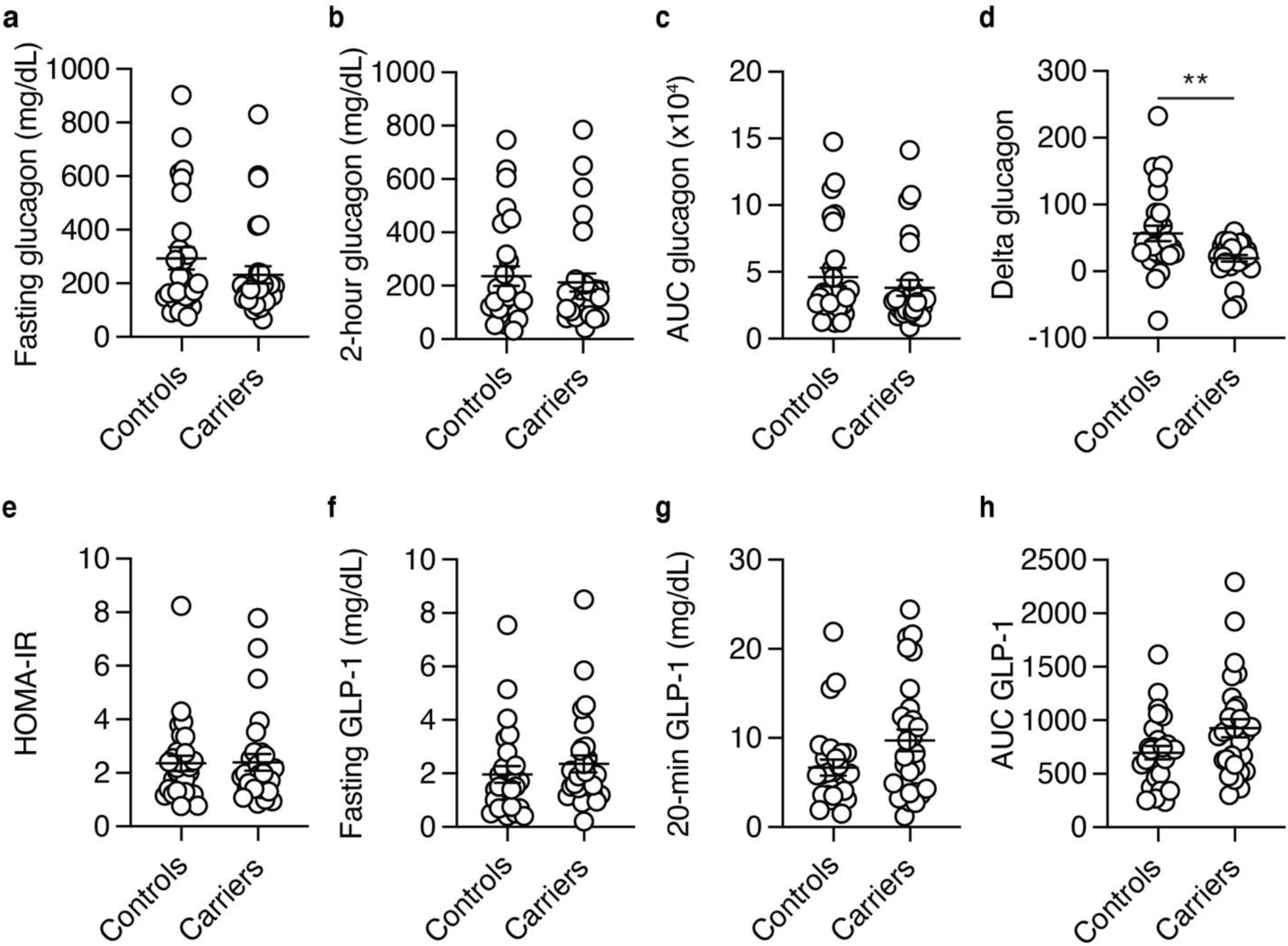
Clinical assessment of glucagon, HOMA-IR, and GLP-1 in carriers of p.Arg192His *PAX4* variant. (**a-d**) Plasma glucagon level (mg/dL) at (**a**) fasting, (**b**) 2-hour time point, (**c**) area under the curve (AUC), and (**d**) delta glucagon during oral glucose tolerance test of p.His192 allele carriers (n=29) and p.Arg192Arg controls (n=28). (**e**) HOMA-IR measurement of p.Arg192Arg controls and p.His192 carriers during the 2-hour oral glucose tolerance test. (**f**) Fasting, (**g**) 20-min, and (**h**) AUC GLP-1 measurements during oral glucose tolerance test. Data are presented as mean±SEM. Statistical analyses were performed using unpaired t-test. *p<0.05, **p<0.01.

**Extended Data Fig. 2.**
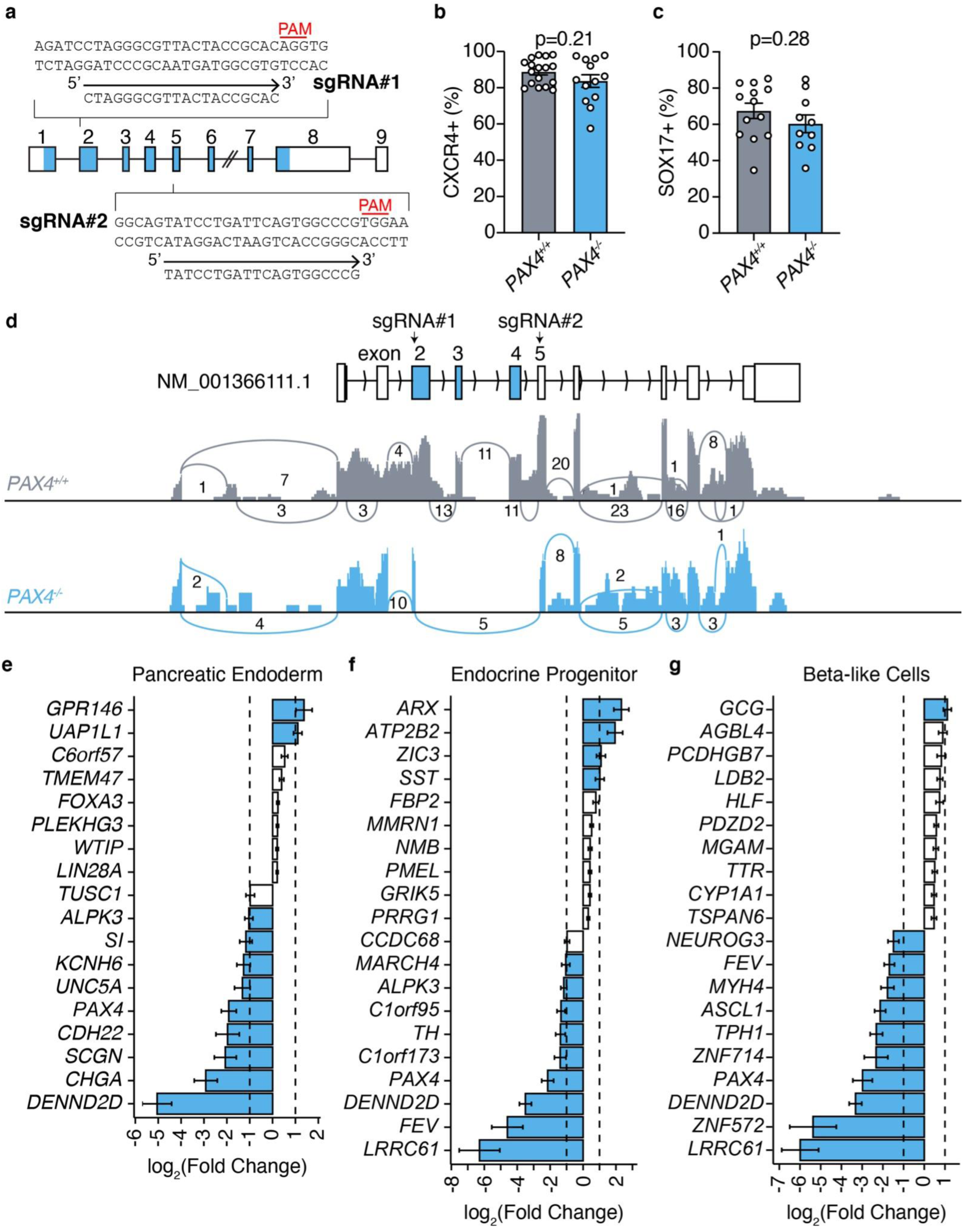
Validation of *PAX4^-/-^* human induced pluripotent stem cells. (**a**) CRISPR-Cas9 genome editing strategy to generate *PAX4*^-/-^ hiPSC isogenic line. Two sgRNAs were designed to target exon 2 (sgRNA#1) and exon 5 (sgRNA#2). PAM genomic sequence is highlighted in red. (**b-c**) Flow cytometry assessment of definitive endoderm markers (**b**) CXCR4 and (**c**) SOX17 of wildtype (*PAX4^+/+^*) and *PAX4*-knockout (*PAX4^-/-^*) DE cells. (**d**) Sashimi plot of *PAX4* transcript from Protocol A confirmed the loss of exons 2 through 5 in *PAX4*^-/-^ lines. (**e-g**) Log_2_(Fold Change) expression of top differentially expressed genes that are expressed in (**e**) pancreatic endoderm, (**f**) endocrine progenitor, and (**g**) beta-like cell stages. Blue bars represent genes with a log_2_(Fold Change) >1 or <-1.

**Extended Data Fig. 3.**
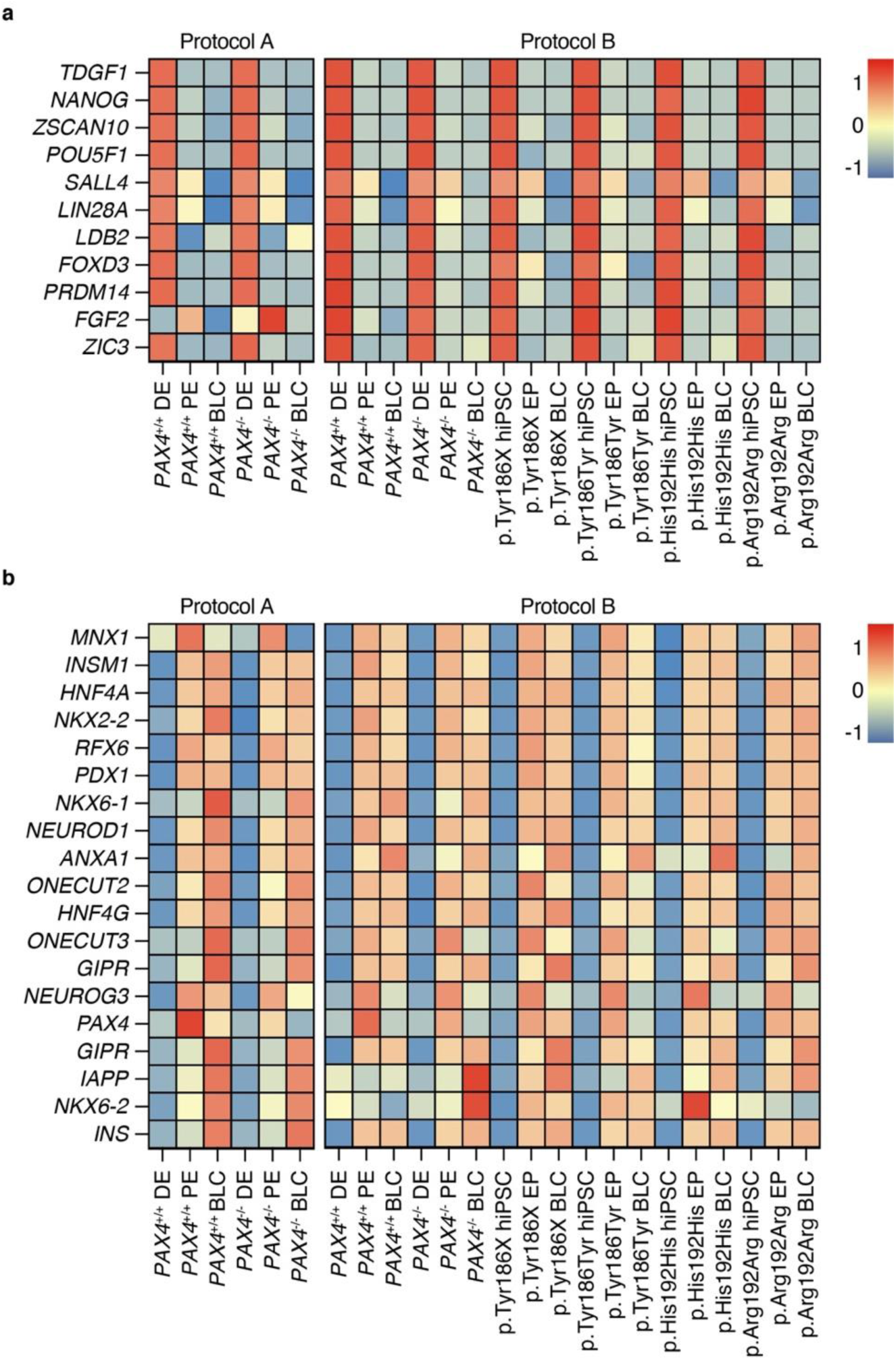
*PAX4*^-/-^ and variant lines have similar repression of pluripotency and activation of endocrine genes as wildtype and corrected lines. (**a**) Key pluripotency and (**b**) endocrine progenitor gene expression in hiPSCs, DE cells, EPs, and BLCs differentiated using Protocols A and B of *PAX4* wildtype (*PAX4*^+/+^), knockout (*PAX4*^-/-^), *PAX4* variants (p.His192His and p.Tyr186X), and corrected (p.Arg192Arg and p.Tyr186Tyr) donor-derived hiPSC lines.

**Extended Data Fig. 4.**
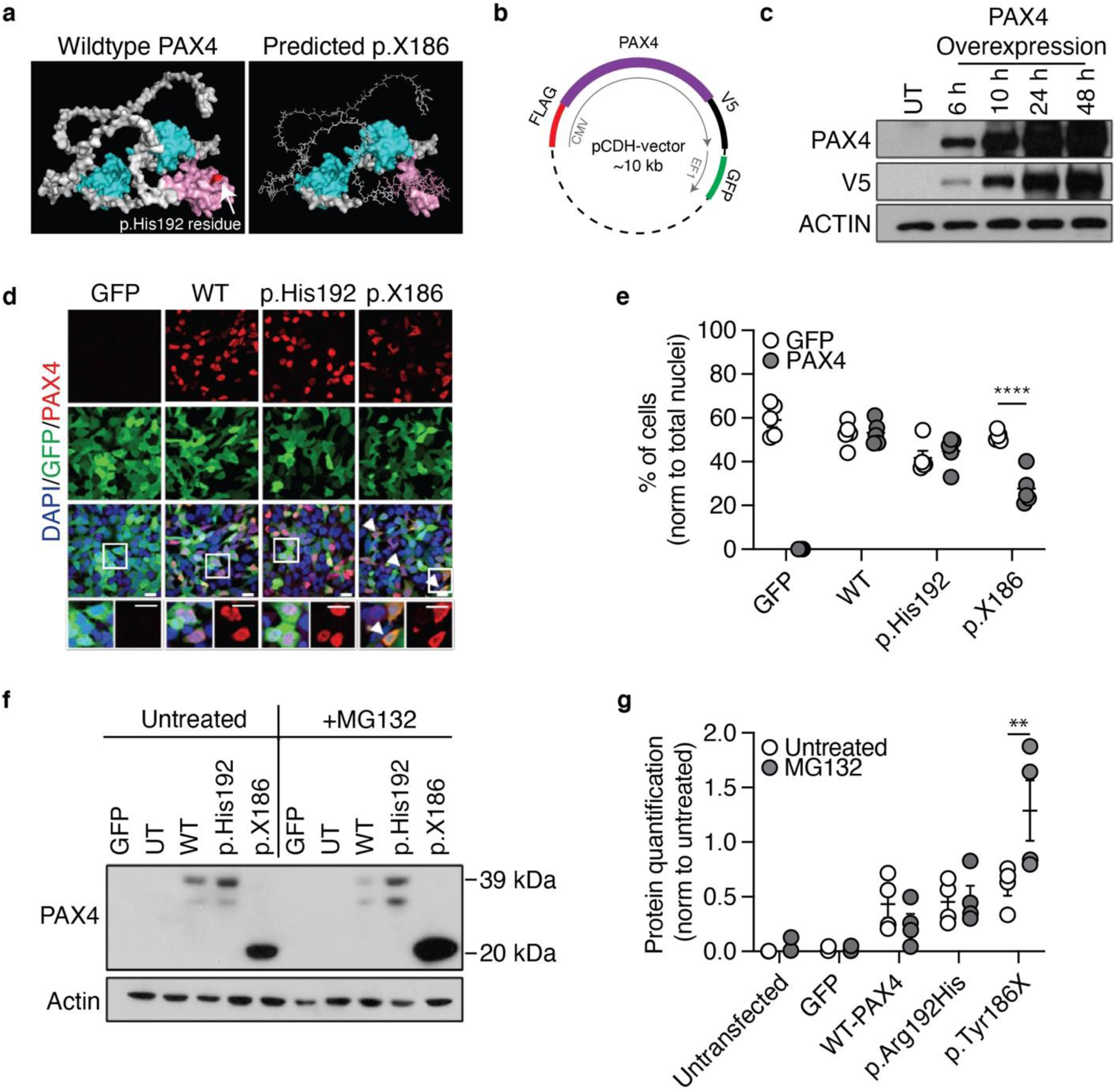
Characterization of PAX4 and its variant proteins. (**a**) Predicted PAX4 protein structure obtained from AlphaFold (AF-O43316-F1-model_v2). PyMOL was used for molecular visualization. Using wildtype PAX4 as template, p.X186 protein was extrapolated to demonstrate protein truncation. (**b**) Construct design for PAX4 overexpression studies. (**c**) Western blot assessment of PAX4 protein, V5 tag (∼37 kD) and ACTIN loading control in AD293 cells transfected with pCDH-WT-PAX4 plasmid for 6, 10, 24 and 48 hours compared to untransfected (UT) control. (**d**) Representative immunofluorescent images of PAX4 (red), GFP (green), and nuclei (DAPI; blue) in AD293 cells following transfection of WT PAX4, p.His192, or p.X186 expressing plasmids. Scale bar = 10 μm. (**e**) Quantification of GFP- and PAX4-expressing cells from immunofluorescence in (**d**). Percentage of cells expressing PAX4 or GFP was normalized to the total number of nuclei (DAPI). Statistical analyses were performed using two-way ANOVA and Sidak’s multiple comparisons test, ****p<0.0001. (**f**) Representative image of western blot assessment and (**g**) densitometry quantification for WT PAX4, p.His192 and p.X186 was overexpressed in AD293 cells and normalized to ACTIN loading control. Cells were treated with or without 10 μM of MG132 for 24 hours posttransfection. Molecular weights of 37 kD and 20 kD correspond to WT PAX4 and p.X186 truncated protein, respectively. n = 4. Statistical analyses were performed using two-way ANOVA and Sidak’s multiple comparisons test, **p<0.01.

**Extended Data Fig. 5.**
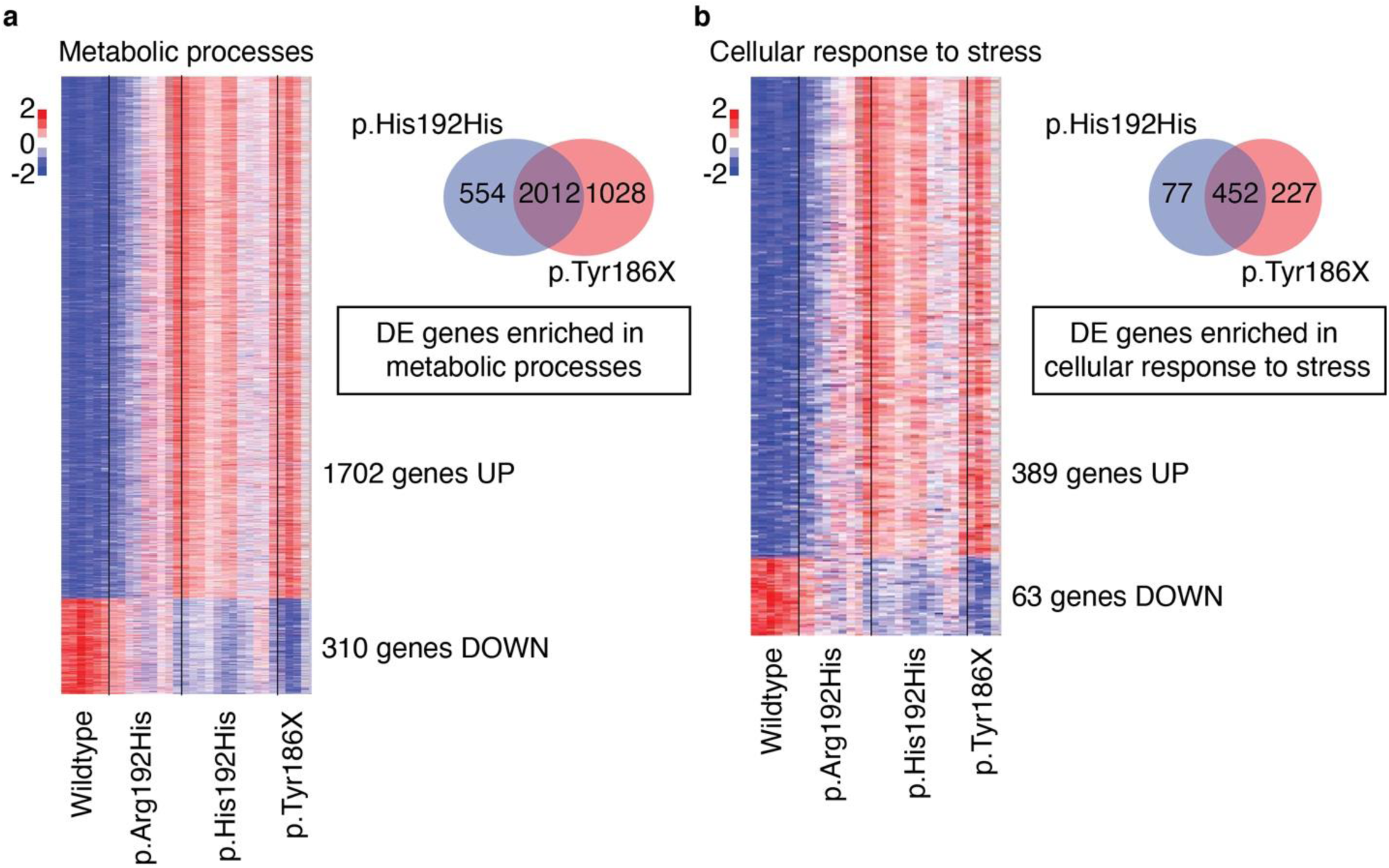
RNA-seq revealed elevated metabolic stress in endocrine progenitors derived from donor-derived hiPSCs carrying *PAX4* variants. Targeted heatmap of differentially expressed genes in endocrine progenitors that are involved in (**a**) metabolic processes (total gene count: 2012; upregulated: 1702; downregulated: 310) and (**b**) cellular response to stress (total gene count: 452; upregulated: 389; downregulated: 63). Venn diagram illustrating differentially expressed (DE) genes enriched in GO terms (**a**) metabolic processes or (**b**) biological processes when comparing p.His192His or p.Tyr186X against *PAX4^+/+^* with FC < 0.5 or FC > 2.

**Extended Data Fig. 6.**
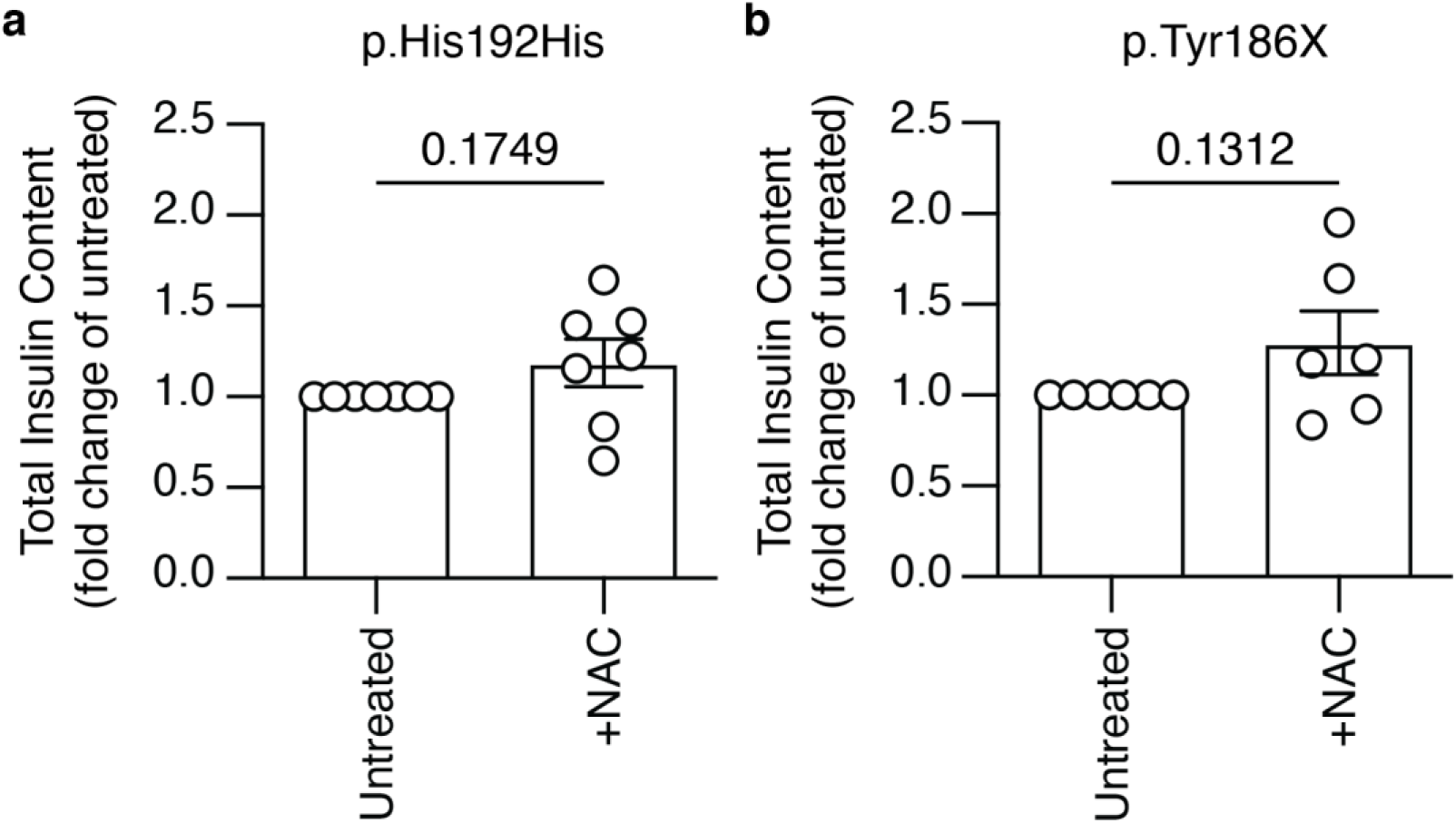
Antioxidant treatment does not rescue the total insulin content in compromised BLCs. Total insulin content of BLCs treated with 10 μM antioxidant NAC from EP to BLC stage carrying (**a**) p.His192His or (**b**) p.Tyr186X. Each dot represents an average of technical replicates of one hiPSC line from one experiment. n>3. Data are presented as mean±SEM. Statistical analyses were performed by Student’s t-test, *p<0.05.

**Extended Data Fig. 7.**
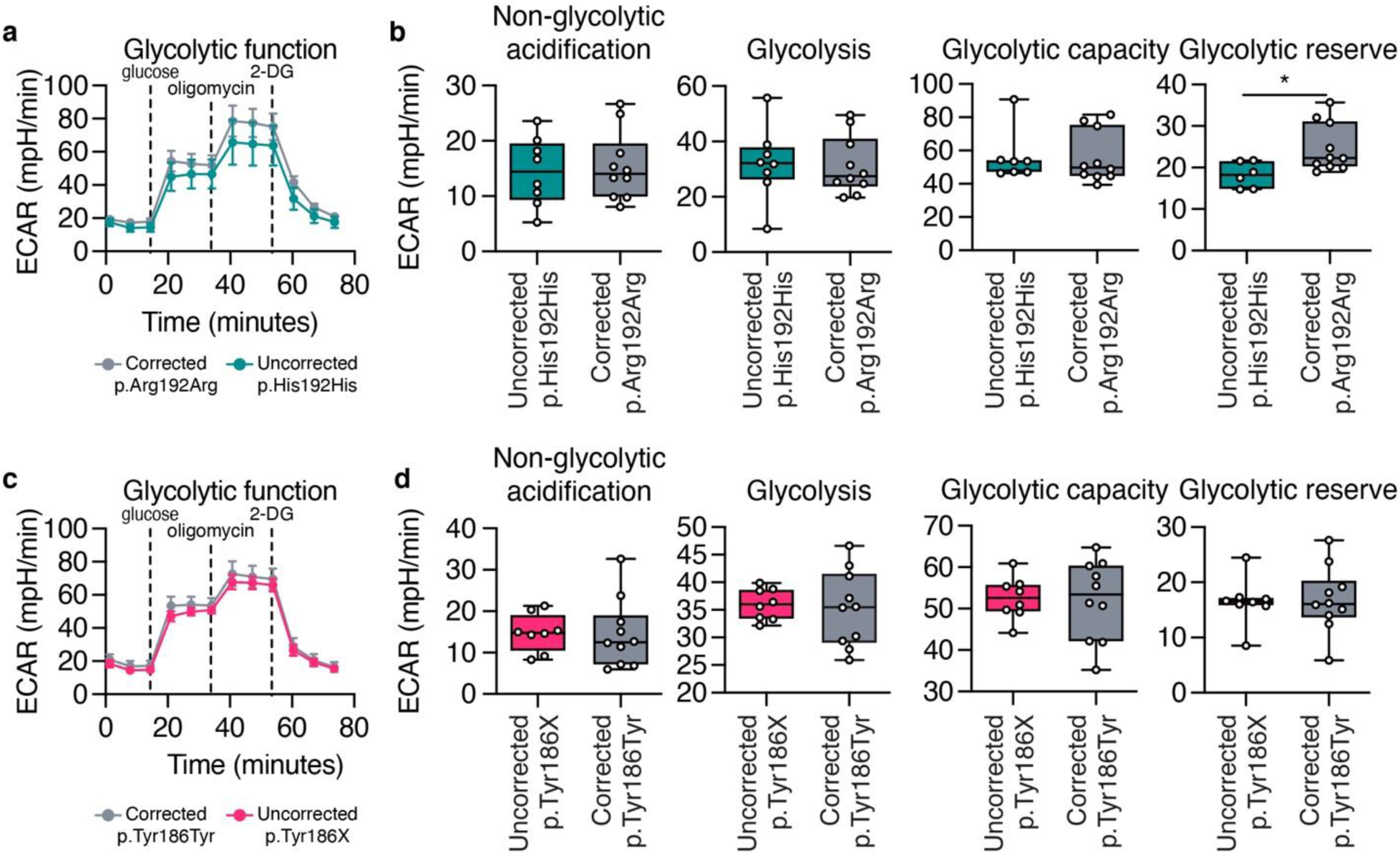
Metabolic stress is not the main causative factor for compromised BLCs. Glycolysis stress test on hiPSC-derived EP cells generated using protocol B. Extracellular acidification rate (ECAR) profiles of EP cells of (**a-b**) p.Arg192Arg (corrected) against p.His192His (uncorrected) and (**c-d**) p.Tyr186Tyr (corrected) against p.Tyr186X (uncorrected). Each data point represents the average measurement rate of technical replicates from one cell line. n>3. Data are presented as mean±SEM. Statistical analyses were performed by Student’s t-test, *p<0.05.

## Supplementary Tables

**Table S1: Aggregated gene-level exome-sequencing association data from Common Metabolic Disease Portal and UKBioBank.**

**Table S2: Differential Expression Analysis of *PAX4*^+/+^ and *PAX4*^-/-^ cells differentiated using Protocol A, Related to** **Figure 3**.

**Table S3: Differential Expression Analysis of *PAX4*^+/+^ and *PAX4*^-/-^ cells differentiated using Protocol B, Related to** **Figure 3**.

**Table S4: Differential Expression Analysis of donor-derived hiPSC lines (*PAX4*^+/+^, p.Arg192His, p.His192His, p.Tyr186X) differentiated using Protocol B, Related to** **Figure 4**.

**Table S5: Differential Expression Analysis of donor-derived hiPSC lines (uncorrected p.His192His and CRISPR-Cas9 corrected p.Arg192Arg) differentiated using Protocol B, Related to** **Figure 7**.

**Table S6: Differential Expression Analysis of donor-derived hiPSC lines (uncorrected p.Tyr186X and CRISPR-Cas9 corrected p.Tyr186Tyr) differentiated using Protocol B, Related to** **Figure 7**.

## Notes

### Competing Interest Statement

Spouse of A.L.G. is an employee of Genentech and holds stock options in Roche. A.K.K.T. is a co-founder of BetaLife Pte Ltd.

