## Supplemental Table 1 for "PAX4 loss of function alters human endocrine cell development and influences diabetes risk"

**Supplementary Table 1a. Gene based association analysis of *PAX4* variants with HbAc1 levels in 268,753 exomes from UKBioBank.**

| **Variant Class** | **Number of Variants** | **P value SKAT-O** | **P Value Burden** | **P Value SKAT** | **CAF** |
| --- | --- | --- | --- | --- | --- |
| pLOF | 20 | 0.0421 | 0.0228 | 0641 | 2.03e-4 |
| Missense | 189 | 0.00683 | 0.0159 | 0.00464 | 0.797 |
| Synonymous | 73 | 0.645 | 0.472 | 0.844 | 0.0757 |

CAF, cumulative allele frequency for this gene (sum of allele frequencies). Data retrieved from [www.genebass.org](http://www.genebass.org) April 2022

**Supplementary Table 1b. Gene based association analysis of *PAX4* variants with type 2 diabetes in 52,000 exomes from the type 2 diabetes knowledge portal.**

| **Test** | **Mask** | **Number of Variants** | **Z score** | **P value** | **Odds Ratio** | **Standard Error** | **Sample Size** |
| --- | --- | --- | --- | --- | --- | --- | --- |
| SKAT | LofTee | 6 | 146476.01 | 0.053 |  |  | 43,125 |
| Collapsing burden |  | 6 | 1.70 | 0.088 | 1.21 | 0.111 | 43,125 |
| Variable threshold |  | 6 | 1.70 | 0.192 | 1.21 | 0.111 | 43,125 |
| SKAT-optimal |  | 6 | 73238.00 | 0.078 |  |  | 43,125 |
| SKAT | 16/16 | 6 | 146476.01 | 0.053 |  |  | 43,125 |
| Collapsing burden |  | 6 | 1.70 | 0.088 | 1.21 | 0.111 | 43,125 |
| Variable threshold |  | 6 | 1.70 | 0.192 | 1.21 | 0.111 | 43,125 |
| SKAT-optimal |  | 6 | 73238.00 | 0.078 |  |  | 43,125 |
| SKAT | 11/11 | 33 | 1093741.78 |  |  |  | 43,125 |
| Collapsing burden |  | 33 | 1.4796 | 0.139 | 1.05 | 0.034 | 43,125 |
| Variable threshold |  | 33 | 2.658 | 0.040 | 1.12 | 0.043 | 43,125 |
| SKAT-optimal |  | 33 | 546870.89 |  |  |  | 43,125 |
| SKAT | **5/5** | 39 | 24368973.87 | **0.00001** |  |  | 43,125 |
| Collapsing burden |  | 39 | 4.34 | **0.000014** | 1.06 | 0.013 | 43,125 |
| Variable threshold |  | 39 | 4.34 | **0.00013** | 1.06 | 0.013 | 43,125 |
| SKAT-optimal |  | 39 | 16026800.08 | **0.00002** |  |  | 43,125 |
| SKAT | **5/5+**  **LofTee LC** | 40 | 24419808.29 | **0.00004** |  |  | 43,125 |
| Collapsing burden |  | 40 | 4.43 | **9.33e-6** | 1.06 | 0.013 | 43,125 |
| Variable threshold |  | 40 | 4.43 | **0.00008** | 1.06 | 0.013 | 43,125 |
| SKAT-optimal |  | 40 | 16530425.89 | **0.00002** |  |  | 43,125 |
| SKAT | **5/5 +1/5 1%** | 77 | 226078120.98 | **1.05e-31** |  |  | 43,125 |
| Collapsing burden |  | 77 | 8.85 | **8.47e-19** | 1.06 | 0.007 | 43,125 |
| Variable threshold |  | 77 | 8.85 | **1.69e1-7** | 1.06 | 0.007 | 43,125 |
| SKAT-optimal |  | 77 | 126954564.88 | **9.26e-31** |  |  | 43,125 |
| SKAT | **5/5 + 0/5 1%** | 92 | 227251674.19 | **5.03e-31** |  |  | 43,125 |
| Collapsing burden |  | 92 | 8.91 | **5.30e-19** | 1.06 | 0.007 | 43,125 |
| Variable threshold |  | 92 | 8.91 | **1.06e-17** | 1.06 | 0.007 | 43,125 |
| SKAT-optimal |  | 92 | 129683675.78 | **2.90e-31** |  |  | 43,125 |

Data retrieved from <https://t2d.hugeamp.org> April 2022. Masks are described in Flannick et al Nature 2018.
